# Reprogramming VHL with molecular glues enables selective degradation of caspase-2

**DOI:** 10.64898/2026.08.12.744529

**Authors:** Jianzhong Hu, Weixian Deng, Shu-Ching Ou, Alex Golkar, Alison Inglis, Kate Smither, Shiqian Li, Kui Chen, Kelvin Cho, Sung Jun Bae, Stephan Zech, Kaylee Choi, Willem den Besten, Stephanie Voss, Olivier Bedel, Bo Zhou, Patrick Ryan Potts, Amine Sadok, Jaeki Min

## Abstract

Molecular glue degraders (MGDs) reprogram E3 ligases to eliminate neosubstrates, yet their application has largely been confined to CRBN. Here, we identify caspase-2 as a new neosubstrate for von Hippel-Lindau (VHL), expanding the scope of VHL-based MGDs. Guided by a focused VHL ligand library design, we employed TurboID-based proximity labeling to discover stereoisomeric compounds (dCASP2-1 and dCASP2-2) that selectively recruit caspase-2 to VHL and promote its ubiquitin-proteasome system-dependent degradation. Further structure-activity relationship (SAR) studies yielded dCASP2-3 and dCASP2-4, which enhanced degradation potency (by 622-fold relative to dCASP2-1) and abolished enantioselectivity. Mechanistic mapping localized the degrader-induced interface to a two-helix region of the caspase-2 CARD domain, with residues H33, P34, and D100 essential for VHL engagement. Degron-guided computational modeling of the VHL/MGD/caspase-2 ternary complex provided structural insight into neosubstrate recognition. Together, we report the development of VHL molecular glues that selectively and potently degrade caspase-2, offering chemical probes to interrogate its functions in apoptosis and stress responses, while broadening the substrate landscape of VHL-based MGDs.

## Introduction

Targeted protein degradation (TPD) has emerged as a powerful therapeutic strategy that utilizes native protein recycling machinery (i.e., ubiquitin-proteasome system, UPS) to remove disease-related proteins^1,2^. By inducing proximity between an E3 ubiquitin ligase and a protein of interest, TPD modalities enable ubiquitination and subsequent degradation of the target protein. There are two major classes of degraders for TPD: proteolysis-targeting chimeras (PROTACs) and molecular glue degraders (MGDs)^3-7^. PROTACs are heterobifunctional molecules composed of two high-affinity protein-binding ligands connected through a linker, while MGDs are typically monovalent small molecules that stabilize or induce a protein-protein interaction between an E3 ligase and a neosubstrate^5^. Although both PROTACs and MGDs induce event-driven target degradation, MGDs may offer better drug-like properties, including lower molecular weight, improved membrane permeability, and greater compliance with Lipinski’s Rule of Five, making them more attractive candidates for clinical development^8^.

Clinically, the MGD class has emerged as a viable and impactful therapeutic modality. Immunomodulatory imide drugs (IMiDs), including thalidomide, lenalidomide, and pomalidomide, are the most established MGDs in the market^9-12^. These agents bind to cereblon (CRBN), the substrate receptor of the CUL4-CRBN E3 ligase complex and reprogram it to engage neosubstrates such as Ikaros (*IKZF1*), Aiolos (*IKZF3*), and CK1α (*CSKN1A1*) for ubiquitination and subsequent proteasomal degradation^10,13,14^. This strategy has shown efficacy in the treatment of multiple myeloma and other hematological malignancies^15^. More recently, a new generation of CRBN-binding MGDs, including CC-92480 (mezigdomide), have shown improved potency and substrate specificity in preclinical and clinical studies^16^, further reinforcing the therapeutic utility of this modality.

Comprehensive proteomic and chemo-proteomic studies have revealed that CRBN recognizes a broad, ligand-programmable spectrum of neosubstrates, including proteins harboring the canonical β-hairpin G-loop degron as well as substrates recruited through emerging structural features such as surface mimicry^17-20^. These findings highlight the adaptability of CRBN for molecular glue discovery and illustrate why CRBN remains the most exploited E3 ligase in clinical-stage MGDs^18,19^. Nevertheless, the dominance of CRBN-based degraders emphasized an unmet need to extend this strategy to other E3 ligases with distinct substrate landscapes and pharmacological profiles.

von Hippel-Lindau (VHL) functions as the substrate recognition subunit of the CUL2-Rbx1 E3 ubiquitin ligase complex and plays a central role in cellular oxygen sensing by selectively recognizing prolyl-hydroxylated HIF-1α hydroxyl proline and promoting its ubiquitination and proteasomal degradation under normoxic conditions^21^. Due to its robust cellular expression, well-defined binding pocket, and the availability of potent, extensively profiled ligands (e.g., VH032, VH298)^22-24^, VHL has been widely employed in PROTAC design and is considered as one of the most versatile E3 ligases for TPD^24-29^. However, despite being widely used for TPD, VHL has been rarely utilized in molecular glue discovery compared to CRBN. To date, only two examples of VHL-based MGDs have been described, which reprogram VHL to degrade the neosubstrates CDO1^30^ and GEMIN3^31^, providing evidence that the VHL ligase can indeed be chemically reprogrammed to engage non-canonical substrates.

Caspase-2 is an evolutionarily conserved, initiator-like protease implicated in differentiation, apoptosis, genome surveillance, and stress responses^32-35^, yet its physiological roles remain debated because activation occurs through the PIDDosome and alternative platforms and yields context-dependent phenotypes^35,36^. Efforts to study its functions have been limited by the lack of selective pharmacological tools since conventional caspase inhibitors display broad cross-reactivity. As a result, caspase-2 biology has been studied primarily through genetic models^35^. This limitation has slowed mechanistic studies, and novel chemical modalities to modulate caspase-2 are highly needed.

In this study, we expanded the scope of VHL-based molecular glues by identifying caspase-2 as a novel VHL neosubstrate. Using proximity-labeling technologies as a target-agnostic screening approach followed up by global proteomics validation, we discovered small molecule VHL ligands that facilitate the assembly of a VHL-MGD-caspase-2 ternary complex, promoting VHL-dependent degradation of endogenous caspase-2. Through degron mapping with domain truncation, motif swapping, and site-directed mutagenesis, we revealed key structural features and interactions that drive caspase-2 recruitment to a degrader-induced interface on VHL. These insights provide a framework for the discovery of VHL-based MGDs and highlight the potential to diversify E3 ligase usage beyond CRBN for therapeutic discovery.

## Results

### TurboID screening identified caspase-2 as a neosubstrate for VHL

To expand the target space for VHL, we developed a state-of-the-art chemo-proteomics screening platform as a target agnostic method to identify novel VHL substrates. We engineered a stable Jurkat cell line that expresses a fusion protein of VHL and TurboID, an engineered biotin ligase that covalently labels proteins in proximity with biotin. Endogenous VHL was knocked out to improve labeling efficiency. In this assay, cells were treated with candidate VHL ligands at 10μM in 96-well format, followed by cell lysis, trypsin digestion, streptavidin pull-down and mass spectrometry analysis to identify protein enriched upon compounds treatment (**Fig. 1A**).

**Figure 1.**
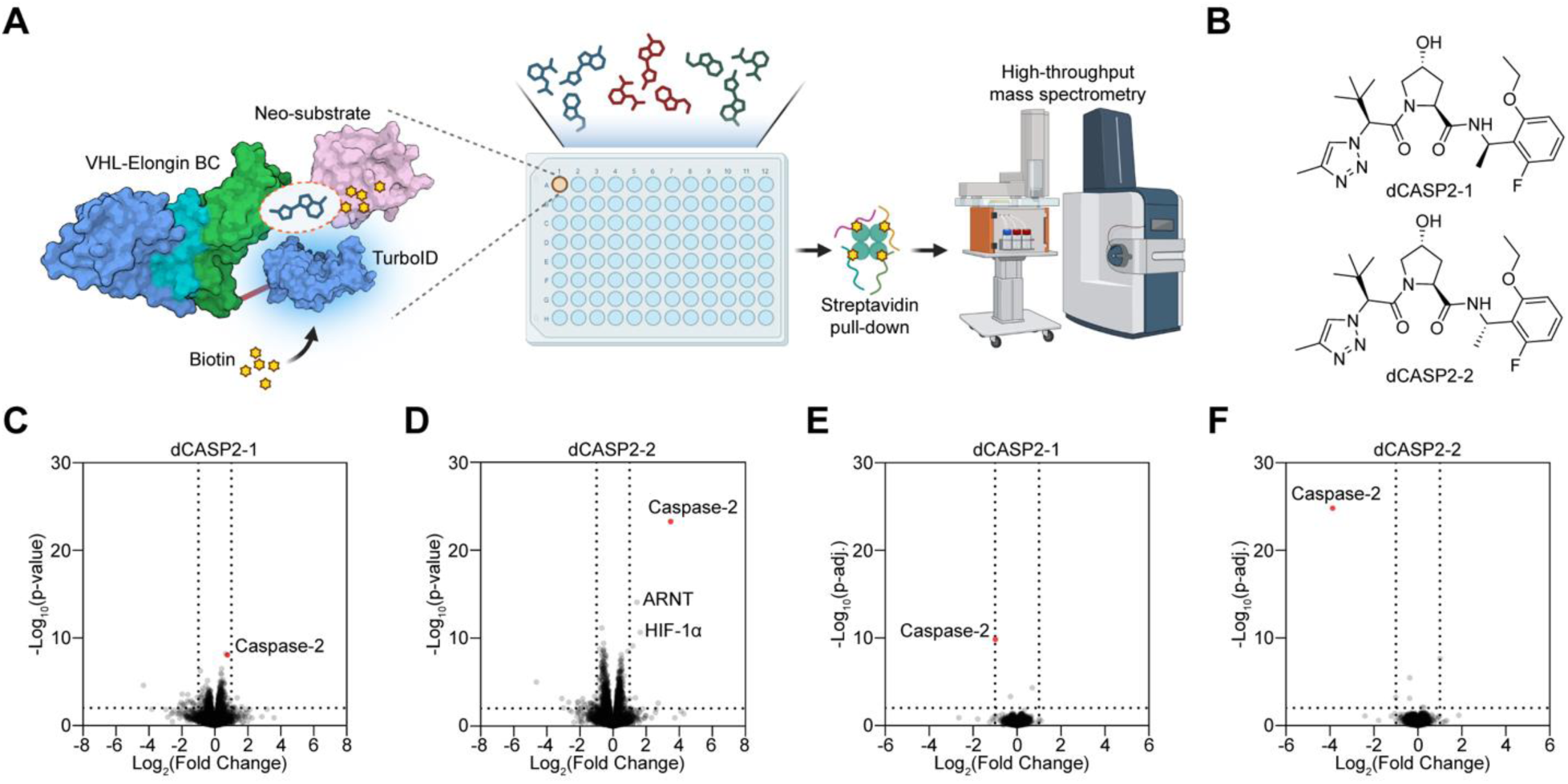
TurboID screening identified caspase-2 as a neosubstrate for VHL. **A.** Diagram of Amgen’s high-throughput TurboID screening platform to discover neosubstrate for VHL. **B.** Chemical structures of dCASP2-1 and dCASP2-2, VHL-binding ligands that recruit caspase-2 to VHL. **C-D.** dCASP2-1 and dCASP2-2 promote proximity between caspase-2 and VHL. Jurkat cells stably expressing TurboID-VHL were treated for 6 hours with bortezomib (2 μM), biotin (50 μM), and either dCASP2-1 (10 μM, **C**, n=3), dCASP2-2 (10 μM, **D**, n=3), or DMSO (0.1%, n=3), followed by streptavidin pull-down and mass spectrometry analysis. Dashed lines indicate significance and enrichment cutoffs of p-value < 0.01 and log_2_ fold-change (relative to DMSO) > 1. Caspase-2 is highlighted in red. **E-F.** dCASP2-1 and dCASP2-2 trigger caspase-2 downregulation. Wild-type Jurkat cells were treated for 24 hours with dCASP2-1 (10 μM, **E**, n=3), dCASP2-2 (10 μM, **F**, n=3), or DMSO (0.1%, n=3), followed by global proteomics analysis. Dashed lines indicate significance and enrichment cutoffs of adjusted p-value < 0.01 and log_2_ fold-change (relative to DMSO) < -1. Caspase-2 is highlighted in red.

Screening of a small, focused VHL ligand library resulted in two compounds, namely dCASP2-1 and dCASP2-2 (**Fig. 1B**) that exhibited activity in selectively inducing proximity between VHL and an unreported neosubstrate caspase-2 (**Fig. 1C-D**). Importantly, dCASP2-1 [*(R)-*] and dCASP2-2 [*(S)-*] are diastereomeric analogs derived from the same chemotype, yet they exhibited a pronounced difference in their ability to recruit caspase-2 to VHL (fold-change of 1.7 for dCASP2-1 versus 11.2 for dCASP2-2, respectively, **Fig. 1C-D**). To validate if the recruitment of caspase-2 to VHL results in degradation, we performed unbiased global proteomics analysis, which confirmed selective degradation of caspase-2 upon 24-h treatment of dCASP2-1 (50% degradation, **Fig. 1E**) and dCASP2-2 (93% degradation, **Fig. 1F**). These results confirmed that dCASP2-1 and dCASP2-2 selectively recruit caspase-2 to VHL, leading to caspase-2 degradation, with the two diastereomers exhibiting differential activity against caspase-2. Interestingly, VHL’s native substrate HIF-1α and its binding partner ARNT (HIF-1β), a reported CRBN neosubstrate^37^, were enriched in the TurboID assay following dCASP2-2 treatment (**Fig. 1D**), despite showing no detectable degradation by global proteomics (**Fig. 1F**). This suggests non-productive induced proximity or a TurboID-specific artifact rather than productive degradation. Although HIF-1α and ARNT have not been reported as enriched hits in VHL-based proximity labeling screens, their enrichment is consistent with the canonical interaction between VHL and HIF-1α and the obligate association of HIF-1α with ARNT.

### VHL ligands dCASP2-1 and dCASP2-2 induce dose-dependent, UPS-mediated caspase-2 degradation

To further validate caspase-2 degradation by these two TurboID hits, we treated Jurkat cells with increasing concentrations of dCASP2-1 or dCASP2-2 for 24 hours, followed by protein lysis and Western blotting. The resulting blots suggest that both dCASP2-1 and dCASP2-2 induced caspase-2 degradation in a dose-response manner (**Fig. 2A-B**). To quantify this effect, we performed JESS analysis to determine levels of caspase-2 degradation after 24-h treatment of dCASP2-1 and dCASP2-2. Both compounds induced concentration-dependent caspase-2 degradation with an estimated D_max_ of 84% for dCASP2-1 and 96% for dCASP2-2, and DC_50_ values of 8.71 μM and 0.051 μM, respectively (**Fig. 2C**, **Table 1, Supplementary Fig. S1**). In parallel, unbiased global proteomics profiling following 6-h compound treatment showed selective loss of caspase-2 by dCASP2-1 and dCASP2-2 (**Supplementary Fig. S2A-B**). This was further validated by Western blotting (**Supplementary Fig. S2C-D**), suggesting the fast-acting nature of these two caspase-2 degraders. Because CDO1 is expressed at low levels in Jurkat cells and may not be reliably captured by TurboID or global proteomics, we directly assessed CDO1 degradation using a HiBiT assay. Our results show neither dCASP2-1 nor dCASP2-2 induced CDO1 degradation (**Supplementary Fig. S2E**), whereas the reported CDO1-targeting degrader VH032 induced robust CDO1 knockdown in RD cells^30^. These results further confirm the selectivity of dCASP2-1 and dCASP2-2 for caspase-2.

**Figure 2.**
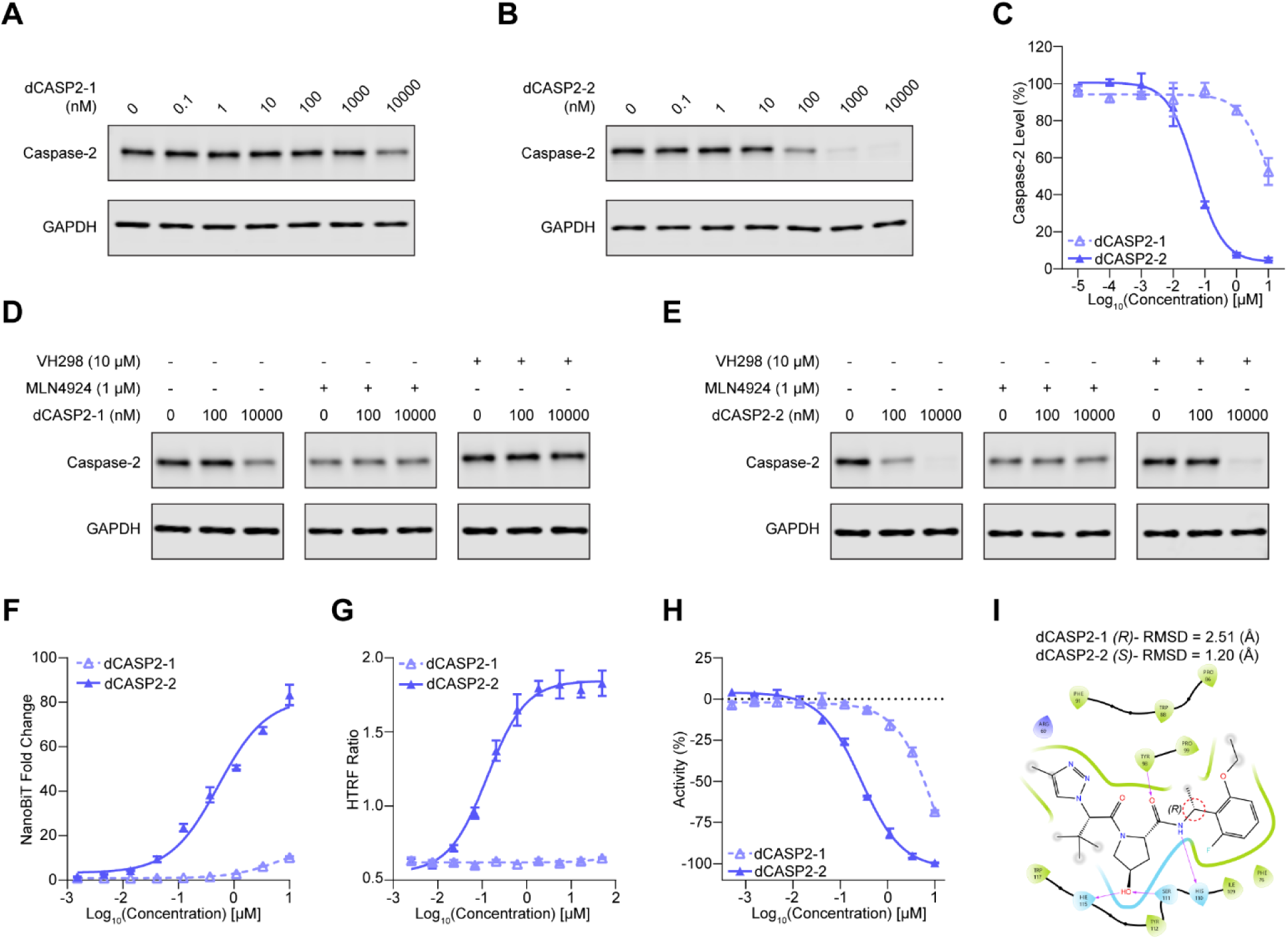
dCASP2-1 and dCASP2-2 mediate UPS-dependent caspase-2 degradation in dose-dependent manner. **A-C.** dCASP2-1 and dCASP2-2 induce caspase-2 degradation in a dose-dependent manner. Jurkat cells were treated for 24 hours with increasing concentrations of dCASP2-1 (**A**) or dCASP2-2 (**B**). Endogenous caspase-2 levels were assessed by Western blot, with GAPDH probed as a loading control. Caspase-2 protein levels were quantified using JESS following 24-hour treatment with dCASP2-1 or dCASP2-2 (**C**, n=3; representative blots are shown in **Supplementary Fig. S1**). **D-E.** Caspase-2 degradation induced by dCASP2-1 and dCASP2-2 is dependent on VHL and the ubiquitin-proteasome system. Jurkat cells were co-treated with increasing concentrations of dCASP2-1 (**D**) or dCASP2-2 (**E**) together with either DMSO, the neddylation inhibitor MLN4924, or the VHL ligand competitor VH298. Caspase-2 levels were analyzed by Western blot. **F.** dCASP2-1 and dCASP2-2 promote caspase-2/MGD/VHL ternary complex formation in cells. HEK293T cells were transiently co-transfected with VHL-LgBiT and SmBiT-caspase-2 vectors, then co-treated with MLN4924 and the indicated degraders. Increased luminescence reflects compound-induced ternary complex formation (n=3). **G.** dCASP2-2 promote *in vitro* caspase-2/MGD/VHL ternary complex formation. HTRF assays were performed with purified caspase-2 and VBC complex in the presence of increasing concentrations of dCASP2-1 or dCASP2-2. Higher HTRF signal indicates ternary complex formation *in vitro* (n=3). **H.** dCASP2-1 and dCASP2-2 bind directly to VHL. VBC AlphaScreen competition assays were used to measure binary binding between the compounds and VHL in vitro. Decreased % activity indicates stronger VHL engagement (n=2). **I.** Computational docking of dCASP2-1 to VHL. The chiral center is highlighted with a red dashed circle. Purple arrows indicate hydrogen bonds. RMSD values for dCASP2-1 and dCASP2-2 are shown at the top.

**Table 1.** Summary of caspase-2 degrader characterization. Fitted D_max_ and DC_50_ values for caspase-2 degradation, VHL engagement values measured by AlphaScreen and permeabilized-cell NanoBRET, and ternary-complex formation (TCF) EC_50_ values measured in cells by NanoBiT and *in vitro* by HTRF are shown. Values were derived from concentration-response curves.

| Compound | dCASP2-1 | dCASP2-2 | dCASP2-3 | dCASP2-4 | VH298 |
| --- | --- | --- | --- | --- | --- |
| **Caspase-2 Degradation**<br>D<sub>max</sub> (%) \| DC<sub>50</sub> (μM) | 83.66 \| 8.71 | 96.31 \| 0.051 | 94.73 \| 0.014 | 95.10 \| 0.014 | 6.13 \| >10 |
| **VHL Engagement EC<sub>50</sub> (μM)**<br>AlphaScreen \| NanoBRET (Perm) | 13.04 \| 5.51 | 0.28 \| 0.39 | 0.23 \| 0.23 | 0.18 \| 0.20 | 0.22 \| 0.59 |
| **Cellular TCF EC<sub>50</sub> (NanoBiT, μM)** | 6.87 | 0.53 | 0.26 | 0.16 | >10 |
| ***In vitro* TCF EC<sub>50</sub> (HTRF, μM)** | >10 | 0.13 | 0.12 | 0.11 | >10 |

To determine whether caspase-2 degradation induced by dCASP2-1 and dCASP2-2 is dependent on the ubiquitin-proteasome system (UPS) and VHL engagement, we performed Western blotting analysis after co-treatment with pathway inhibitors. Our results show co-treatment with MLN4924, a neddylation inhibitor that blocks Cullin-RING E3 ligase activity, fully rescued caspase-2 levels, indicating that degradation requires functional Cullin-RING E3 ligases (**Fig. 2D-E**). In addition, competition with a non-caspase-2 degrading VHL ligand VH298 abrogated caspase-2 degradation (**Fig. 2D-E, Supplementary Fig. S3A**), demonstrating that both caspase-2 degraders act through direct VHL recruitment. Taken together, these results confirm that dCASP2-1 and dCASP2-2 promote UPS-dependent, VHL-mediated degradation of caspase-2.

### Ternary complex formation (TCF) between the neosubstrate, degrader, and E3 ligase is required for functional targeted protein degradation

To test whether dCASP2-1 and dCASP2-2 promote caspase-2-VHL engagement in live cells, we employed a NanoBiT complementation assay. Both compounds induced dose-dependent TCF between caspase-2 and VHL (**Fig. 2F**) while VH298 failed to induce cellular TCF (**Supplementary Fig. S3B**), consistent with their ability to induce caspase-2-VHL proximity and caspase-2 degradation (**Fig. 2C, Supplementary Fig. S3A**). Notably, dCASP2-2 elicited stronger NanoBiT signals at lower concentrations compared to dCASP2-1, indicating that it is both more potent and more efficacious in promoting caspase-2-VHL complex formation. To further validate these findings in a biochemical context, we performed an *in vitro* homogeneous time resolved fluorescence (HTRF) assay, which similarly showed degrader-induced TCF between recombinant caspase-2 and VHL (**Fig. 2G, Supplementary Fig. S3C**). The high concordance between the cellular and biochemical assays demonstrates that the observed ternary complex formation is directly mediated by these caspase-2 degraders. Together, these results support a model where enhanced ternary complex formation supports the superior degradation activity of dCASP2-2.

We then quantitatively assessed the binary binding of the two degraders to VHL. We first employed an AlphaScreen assay and found dCASP2-1 exhibited ∼10-fold weaker binding compared to dCASP2-2 (**Fig. 2H**), consistent with their differential degradation activities (**Fig. 2C**). We further evaluated VHL engagement in a cellular context using a NanoBRET displacement assay in both live and permeabilized cells. In both settings, dCASP2-2 exhibited substantially greater displacement of the tracer compared to dCASP2-1 (**Supplementary Fig. S2F-G**), indicating that the difference in degradation activity arises from differential direct VHL binding rather than variations in cellular permeability. Notably, the binding affinity of dCASP2-2 for VHL was comparable to that of the well-characterized high affinity VHL binder VH298 (**Supplementary Fig. S3D-F**), indicating its potent engagement of the VHL binding pocket. To further probe the structural basis of the differential VHL affinity between the two diastereomers, we performed computational docking of both compounds into the VHL binding pocket. The analysis indicated that dCASP2-2 adopts a binding pose with a lower root-mean-square deviation (RMSD) relative to dCASP2-1 (**Fig. 2I**), suggesting that a more favorable binding conformation may contribute to enhanced VHL binding and degradation activity.

### Chemical modification of VHL ligand RHS abolishes enantioselective caspase-2 degradation

Given the pronounced stereoselectivity observed between dCASP2-1 [*(R)-*] and dCASP2-2 [*(S)-*], we investigated whether structural modifications could enhance VHL binding affinity and caspase-2 degradation activity while attenuating stereochemical preference. To this end, we modified the right-hand side (RHS) of the moiety by replacing the ethoxy-substituted aromatic group with a fused cyclic ether, dihydrobenzopyran, to reduce conformational flexibility while preserving the VHL-binding motif. This strategy afforded two new analogs, dCASP2-3 [*(R)-*] and dCASP2-4 [*(S)-*], both incorporating the modified RHS while retaining the VHL-binding core (**Fig. 3A**). To test if such structural modification could attenuate the stereoselectivity observed between dCASP2-1 and dCASP2-2, we tested caspase-2 degradation followed by 24-h treatment of dCASP2-3 or dCASP2-4 by Western blotting. We observed similar level of caspase-2 degradation for both compounds (**Fig. 3B-C**). This finding was further confirmed by quantitative JESS analysis, which showed that both dCASP2-3 and dCASP2-4 elicited similar extents of caspase-2 degradation activity, with both compounds achieving a Dmax of 95% and a DC50 of 0.014 μM (**Fig. 3D**, **Table 1, Supplementary Fig. S1**). Consistent with this, co-treatment with MLN4924 or VH298 rescued caspase-2 levels (**Fig. 3E-F**), confirming UPS- and VHL-dependent degradation by both degraders.

**Figure 3.**
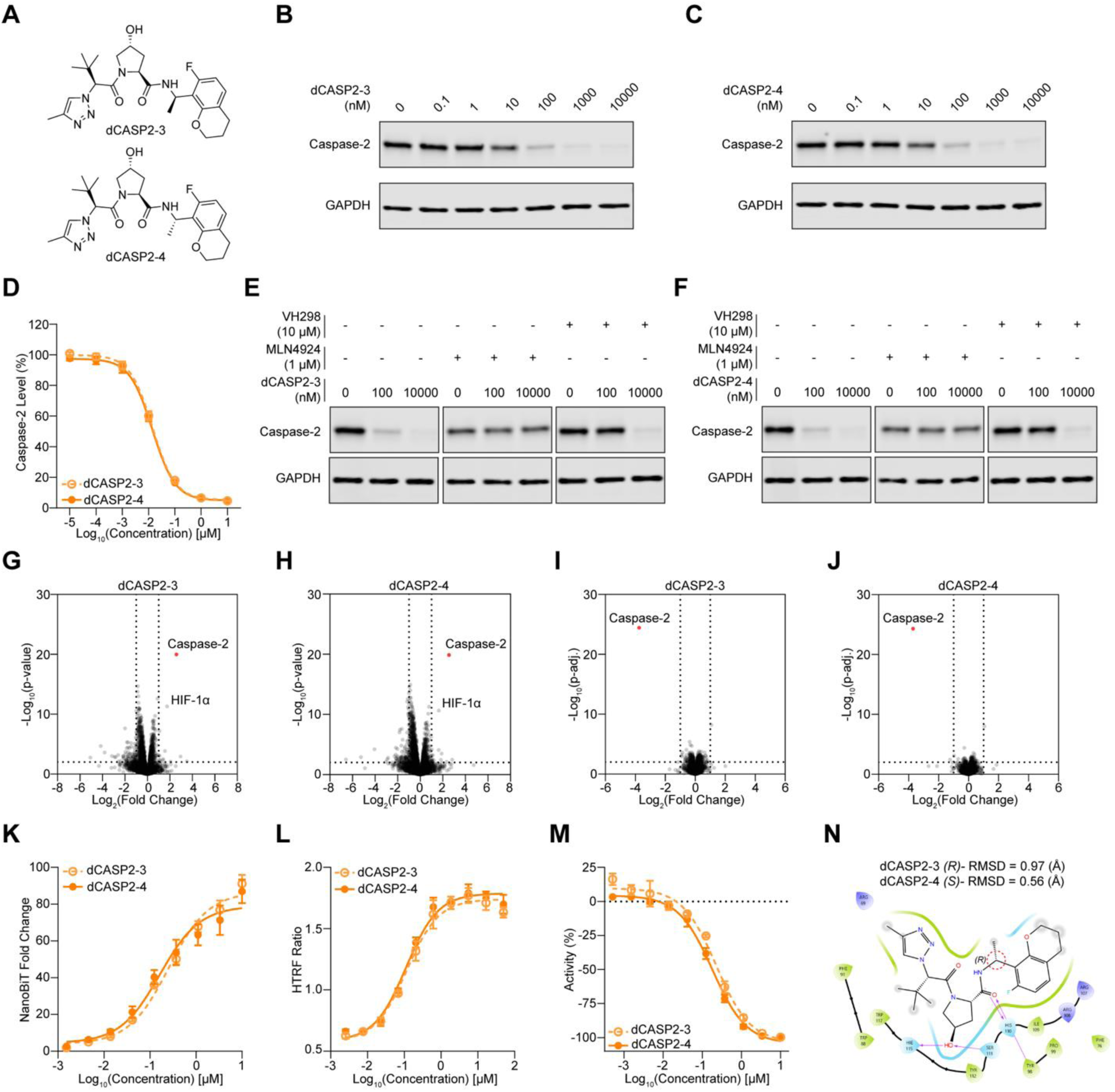
Chemical modification of VHL ligand RHS abolished enantioselectivity for VHL binding and Caspase2 degradation. **A.** Chemical structures of dCASP2-3 and dCASP2-4. **B-D.** dCASP2-3 and dCASP2-4 degrade caspase-2 in a dose-dependent manner. Jurkat cells were treated for 24 hours with increasing concentrations of dCASP2-3 (**B**) or dCASP2-4 (**C**). Endogenous caspase-2 levels were visualized by Western blot, with GAPDH probed as loading control. Caspase-2 protein levels were quantified using JESS following 24-hour treatment with dCASP2-3 or dCASP2-4 (**D**, n=3; representative blots shown in **Supplementary Fig. S1**). **E-F.** Caspase-2 degradation induced by dCASP2-3 and dCASP2-4 is dependent on VHL and the ubiquitin-proteasome system. Jurkat cells were co-treated with increasing concentrations of dCASP2-3 (**E**) or dCASP2-4 (**F**) together with either DMSO, the neddylation inhibitor MLN4924, or the VHL ligand competitor VH298. Caspase-2 levels were analyzed by Western blot. **G-H.** dCASP2-3 and dCASP2-4 promote proximity between caspase-2 and VHL. TurboID-VHL-expressing Jurkat cells were treated for 6 hours with bortezomib (2 μM), biotin (50 μM), and either dCASP2-3 (10 μM, **G**, n=3), dCASP2-4 (10 μM, **H**, n=3), or DMSO (0.1%, n=3). Streptavidin pull-down followed by mass spectrometry was performed. Dashed lines indicate significance and enrichment cutoffs of p-value < 0.01 and log_2_ fold-change (relative to DMSO) > 1. Caspase-2 is highlighted in red. **I-J.** dCASP2-3 and dCASP2-4 induce caspase-2 downregulation. Wild-type Jurkat cells were treated for 24 hours with dCASP2-3 (10 μM, **I**, n=3), dCASP2-4 (10 μM, **J**, n=3), or DMSO (0.1%, n=3), followed by global proteomics analysis. Dashed lines indicate significance and enrichment cutoffs of adjusted p-value < 0.01 and log_2_ fold-change (relative to DMSO) < -1. Caspase-2 is highlighted in red. **K.** dCASP2-3 and dCASP2-4 promote caspase-2/MGD/VHL ternary complex formation in cells. HEK293T cells were transiently co-transfected with VHL-LgBiT and SmBiT-caspase-2 vectors, then co-treated with MLN4924 and the indicated degraders. Increased luminescence reflects compound-induced ternary complex formation (n=3). **L.** dCASP2-3 and dCASP2-4 promote *in vitro* caspase-2/MGD/VHL ternary complex formation. HTRF assays were performed with purified caspase-2 and VBC complex in the presence of increasing concentrations of dCASP2-3 or dCASP2-4. Higher HTRF signal indicates ternary complex formation *in vitro* (n=3). **M.** dCASP2-3 and dCASP2-4 bind directly to VHL. VBC AlphaScreen competition assays were performed to measure binary binding of the compounds to VHL *in vitro*. Decreased % activity indicates stronger VHL engagement (n=2). **N.** Computational docking of dCASP2-3 to VHL. The chiral center is highlighted with a red dashed circle, and purple arrows indicate hydrogen bonds. RMSD values for dCASP2-3 and dCASP2-4 are shown at the top.

To further validate these findings, we performed TurboID proximity labeling upon dCASP2-3 or dCASP2-4 treatment. Both dCASP2-3 and dCASP2-4 exhibited comparable selective recruitment of caspase-2 enrichment to VHL, with approximately 6-fold enrichment observed for dCASP2-3 (**Fig. 3G**) and dCASP2-4 (**Fig. 3H**). The induced enrichment of caspase-2 to VHL was then validated by global proteomics analysis after 24-h treatment of dCASP2-3 (93% degradation, **Fig. 3I**) or dCASP2-4 (92% degradation, **Fig. 3J**). In addition, we tested whether dCASP2-3 or dCASP2-4 can also induce rapid caspase-2 degradation by conducting 6-h proteomic analysis. Notably, both compounds induced caspase-2 degradation after 6-h treatment (**Supplementary Fig. S4A-B**), which was confirmed by Western blotting (**Supplementary Fig. S4C-D**). Importantly, we observed no degradation of CDO1 by either compound using a HiBiT assay (**Supplementary Fig. S4E**).

Finally, we evaluated ternary complex formation and VHL binding. Both compounds promoted caspase-2-VHL engagement in live cells (NanoBiT) and *in vitro* (HTRF) with comparable potency (**Fig. 3K-L**). Binary VHL binding affinities measured by AlphaScreen and NanoBRET (in both live and permeabilized cells) were also similar (**Fig. 3M, Supplementary Fig. S4F-G**). Computational docking supported these observations, revealing similar binding poses and RMSD values for dCASP2-3 and dCASP2-4 (**Fig. 3N**).

Taken together, these data demonstrate that modifying the RHS of the caspase-2 MGDs abolishes the strong enantioselectivity observed with dCASP2-1 and dCASP2-2, yielding stereoisomers with equivalent activity across degradation, ternary complex formation, and VHL binding while maintaining high selectivity against caspase-2. This finding suggests RHS optimization can be leveraged to tune stereochemical dependence in VHL-based MGDs.

### Degron mapping reveals CARD domain as the VHL-interacting element of caspase-2

To identify the caspase-2 region responsible for molecular glue-induced VHL engagement, we employed dCASP2-3 as a tool compound and generated constructs encoding the full-length protein, the CARD domain, or the p19 and p12 subunits (**Fig. 4A**). Using a NanoBRET ternary complex formation (TCF) assay, we observed robust, concentration-dependent TCF only with the full length and isolated CARD domain, whereas the p19 and p12 fragments remained silent (**Fig. 4B**). Thus, the interaction localizes to the CARD domain, which is sufficient for dCASP2-3-induced VHL recruitment.

**Figure 4.**
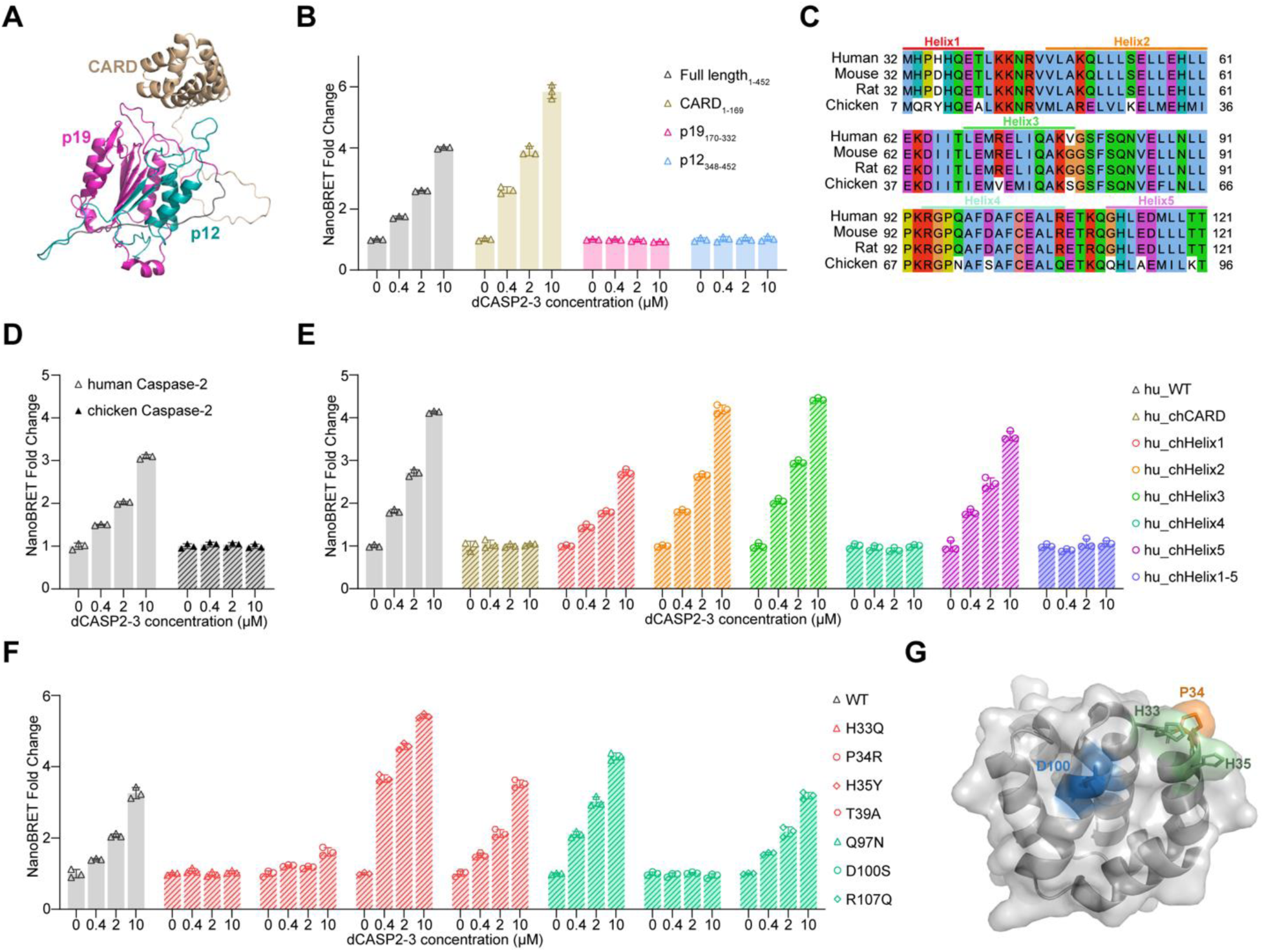
Degron mapping indicates CARD domain interacts with VHL upon dCASP2-3 treatment. **A.** AlphaFold structure of caspase-2. CARD, p19, and p12 domains are labeled in different colors. **B.** NanoBRET ternary complex assays demonstrate that the CARD domain is sufficient for caspase-2/dCASP2-3/VHL interaction. NanoLuc-fused full-length caspase-2 (gray bars) or truncated caspase-2 fragments (colored bars) were co-expressed with Halo-fused VHL in HEK293T cells, followed by co-treatment with MLN4924 (1 μM) and increasing concentrations of dCASP2-3. NanoBRET ratios (618 nm/460 nm) were normalized to DMSO controls (n=3). Increased NanoBRET signal indicates compound-promoted proximity between caspase-2 fragments and VHL. **C.** Sequence alignment of human and chicken caspase-2 CARD domains. **D.** dCASP2-3 fails to recruit chicken caspase-2 to VHL. As described in (**B**), NanoBRET assays were performed using constructs encoding full-length human (gray bars without texture) or chicken (gray bars with strips) caspase-2 (n=3). Increased NanoBRET signal reflects compound-promoted proximity between caspase-2 and VHL. **E.** Helices 1 and 4 within the caspase-2 CARD domain are necessary for VHL interaction. NanoBRET assays were performed with NanoLuc-fused human caspase-2 variants carrying chicken-derived motif substitutions (striped bars indicate constructs contain chicken elements, n=3). **F.** Amino acid residues in Helices 1 and 4 modulate VHL interaction. NanoBRET assays were performed with NanoLuc-fused human caspase-2 variants carrying chicken-derived amino acid substitutions (red bars: substitutions within Helix1, cyan bars: substitutions within Helix4, n=3). **G.** Structure of CARD domain, key residues for VHL engagement are labeled with colors. Blue: acidic residue, green: basic residue, orange: hydrophobic residue.

Sequence alignment of the caspase-2 CARD domain across species showed that the human, mouse, and rat sequences are highly conserved, whereas the chicken CARD domain is more divergent from the human sequence (**Fig. 4C**). We next compared dCASP2-3-induced TCF with human and chicken caspase-2 constructs. The results show dCASP2-3 can only form TCF with human caspase-2 but not with the chicken ortholog (**Fig. 4D**). Although the overall CARD domain fold is conserved, multiple side-chain disparities exist in five out six helices within the CARD domain (**Fig. 4C**), suggesting that the sequence differences in the CARD domain govern VHL engagement.

We next generated a panel of human-chicken chimeras in which individual helices of the human CARD domain were substituted with their chicken derivatives. Swapping human Helix 1 or Helix 4 with their respective chicken counterparts markedly impaired compound-induced TCF, while replacement of other helices had minimal effect (**Fig. 4E**), suggesting helices 1 and 4 are critical interaction elements. We then performed systematic mutagenesis of ortholog-divergent residues within these helices. In the cellular TCF assay, H33Q and P34R in Helix 1 and D100S in Helix 4 abolished ternary-complex formation, whereas H35Y in Helix 1 enhanced the interaction (**Fig. 4F**).

Together, these results define a composite two-helix interface within the caspase-2 CARD domain as the MGD-dependent VHL interaction site (**Fig. 4G**). Specific residues within helices 1 and 4 (H33, P34, D100) act as essential components for neosubstrate recognition, while others (e.g., H35) may further modulate the efficiency of ternary complex formation.

### Degron-informed computational modeling proposes a VHL-MGD-caspase-2 ternary complex

To explore possible molecular mechanism underlying caspase-2 recruitment by dCASP2, we implemented an integrated computational workflow to generate a VHL-MGD-caspase-2 ternary complex, guided by degron-mapping results (**Fig. 4G**). The computation model localized the critical recruitment interface to the caspase-2 CARD domain, with mutational scanning identifying H33, P34, H35, and D100 as key functional residues. These experimental insights informed both the docking restraints and post-docking filters applied throughout the computational pipeline. Because the resulting models were derived by these experimental constraints, they represent structural hypotheses that are consistent with the available data rather than independently determined ternary-complex structures.

We first modeled the VHL-dCASP2-3 binary complex using Glide docking into the VHL structure (PDB: 9BOL)^38^ by enforcing canonical VHL hydrogen bonds with Y98, H110, S111, and H115. We then calculated and compared the local backbone RMSD of the relaxed VHL-dCASP2-3 binary complex and the apo VHL (1VCB)^39^. Superposition of the binary model with apo VHL revealed a highly conserved overall conformation with RMSD difference of 0.8 Å (**Fig. 5A**), indicating that dCASP2-3 binding does not induce major structural rearrangements in VHL.

**Figure 5.**
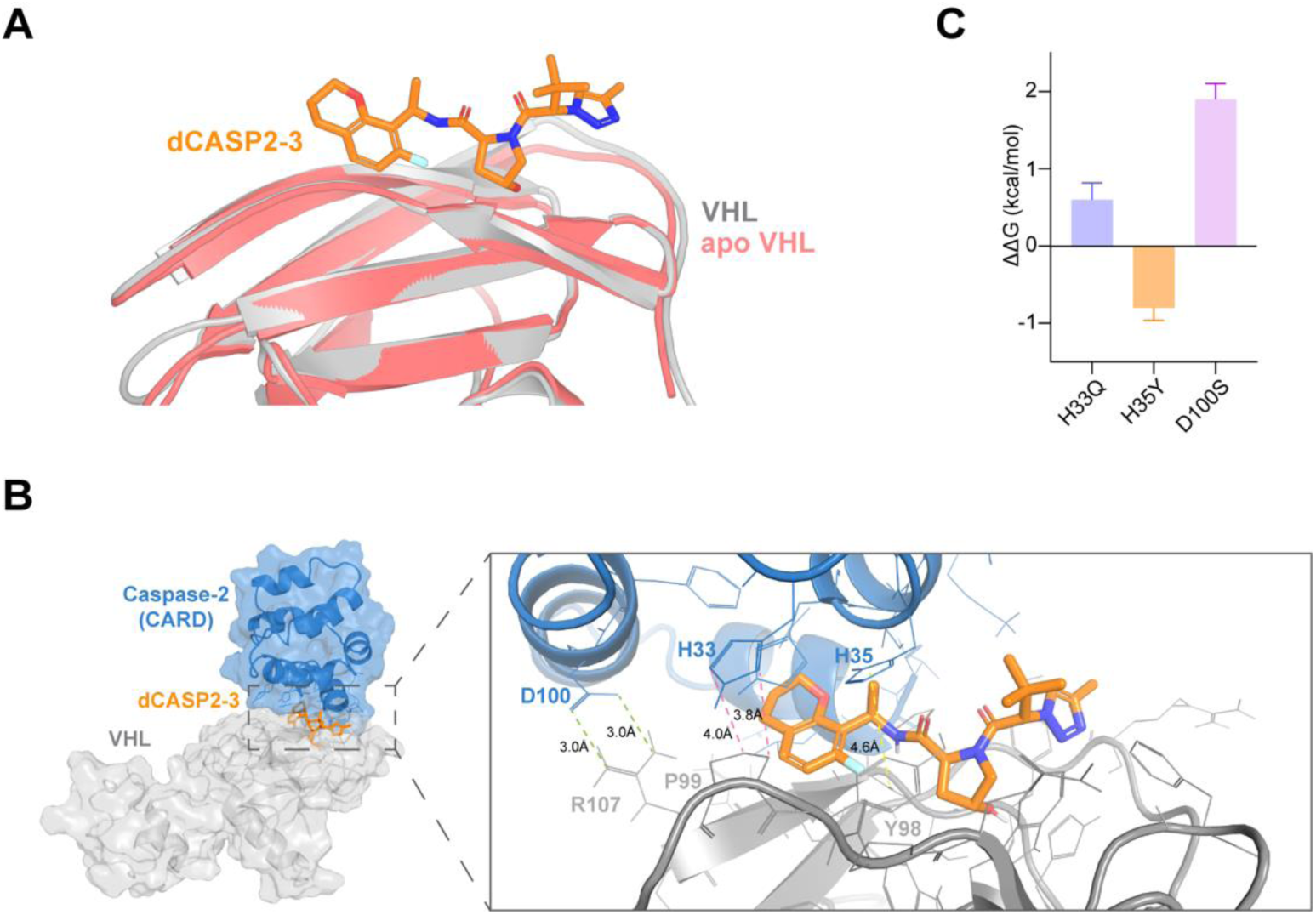
Degron-guided computational model of the caspase-2/dCASP2-3/VHL ternary complex. **A.** Structural comparison of the modeled VHL-dCASP2-3 binary complex and apo VHL. The relaxed dCASP2-3-bound VHL model is shown in gray, apo VHL (PDB 1VCB) in magenta, and dCASP2-3 in orange. Superposition of the two VHL structures yielded a local backbone RMSD difference of 0.8 Å. **B.** Selected model of VHL/dCASP2-3/caspase-2 ternary complex. Caspase-2 CARD domain is shown in blue, VHL in gray, and dCASP2-3 in orange. The bottom panel highlights the ternary interface, with key interactions between caspase-2 (H33, H35, D100) and VHL (Y98, P99, R107) indicated. Green lines: salt bridges; pink lines: hydrophobic packing; yellow lines: π-π aromatic stacking. **C.** Relative binding free energy changes (ΔΔG) for three caspase-2 mutations calculated using Schrödinger’s FEP+ workflow. The uncertainty was determined by MBAR (Multistate Bennett Acceptance Ratio) to estimate the noise induced from sampling.

Next, ternary complex assembly was performed by docking VHL-dCASP2 models to the caspase-2 CARD using PIPER^40^, with experimentally validated degron residues specified as attractive restraints. From 1,000 candidates, we refined 10-15 representative complexes by extended MD. The selected model maintained a coherent VHL-caspase-2 interface with RMSD ≤ 3 Å over 240 ns, consistent with a robust ternary architecture. In this selected model, interaction analysis revealed that dCASP2-3 contributes minimally to direct caspase-2 contacts (**Fig. 5B**). At the modeled VHL-caspase-2 interface, caspase-2 D100 was predicted to form a salt bridge and hydrogen bond with VHL R107 (**Fig. 5B**). We also observed and a H33 (caspase-2)-P99 (VHL) hydrophobic contact and a possible H35 (caspase-2)-Y98 (VHL) π-π interactions (**Fig. 5B**).

To examine whether the selected model was compatible with the degron-mapping data, we used Schrödinger FEP+ to calculate free energy change (ΔΔG) for selected substitutions (H33Q, H35Y, D100S) on caspase-2. The P34R substitution did not converge under the available sampling protocol and was therefore excluded from quantitative analysis. H33Q and D100S were predicted to be destabilizing, whereas H35Y was predicted to promote ternary formation, in directional agreement with the experimental results (**Fig. 5C**). The H35Y model suggested the caspase-2 H35Y sidechain substitution might provide stronger π-π stacking with Y98 (VHL) and additional hydrogen bond with P97 backbone (VHL) (**Supplementary Fig. S5**). These computational outcomes support the functional degron mapping and highlight the centrality of D100 (caspase-2)-R107 (VHL), H33 (caspase-2)-P99 (VHL) and H35 (caspase-2)-Y98 (VHL) interactions to complex stability.

Together, the degron-informed modeling established a plausible mechanistic framework in which dCASP2-3 promotes selective recruitment of caspase-2 by stabilizing interactions between VHL and a specific CARD-based degron. The integrated experimental-computational approach rationalized SAR trends, validated mutational effects, and provided a structural rationale for the selective neosubstrate recognition. However, alternative ternary-complex geometries may also be compatible with the current experimental data, and direct structural studies are required to establish a precise ternary complex model.

## Discussion

Our data established caspase-2 as a novel neosubstrate for the VHL E3 ligase, expanding the target space of VHL-based molecular glue degraders beyond previously reported neosubstrates such as CDO1 and GEMIN3^30,31^. Through a target-agnostic TurboID screening strategy, followed by orthogonal validation with proteomics, Western blotting, HTRF, and quantitative degradation assays, we discovered a series of small molecules (dCASP2-1 to dCASP2-4) that recruit caspase-2 to VHL and trigger its selective degradation in UPS- and VHL-dependent manner. Mechanistic analyses demonstrated that ternary complex formation is the key determinant of degrader potency, with stereochemistry and scaffold modifications modulating both VHL affinity and caspase-2 recruitment efficiency. Notably, a rapid modification of the right-hand side of the degrader scaffold eliminated the strong stereoselectivity observed between dCASP2-1 and dCASP2-2, yielding the stereoisomers dCASP2-3 and dCASP2-4, which exhibit enhanced degradation activity while retaining selectivity for caspase-2. We hypothesize that this effect arises from the role of the RHS substituent in modulating ligand conformation and VHL presentation rather than directly engaging the neosubstrate. In this context, cyclization could constrain the ligand into a more favorable binding geometry confirmed by docking model (**Fig. 2I** for dCASP2-1 and-2 vs **Fig.3N** for dCASP2-3 and - 4), thereby facilitating productive ternary complex formation in a manner that reduces sensitivity to stereochemistry.

By incorporating truncation, ortholog swapping, and systematic mutagenesis, we further mapped the caspase-2 degron to a two-helix region within the CARD domain. Key residues (H33, P34, and D100) were found to be essential for VHL engagement, while H35 positively modulated interaction strength (**Fig. 4F**). Computational modeling further provided a structural hypothesis of VHL/MGD/caspase-2 ternary complex. The selected model was consistent with the observed mutational effects and suggested possible interactions involving caspase-2 D100, H33, and H35 at the VHL interface (**Fig. 5B**). Because the degron-mapping and mutational data were used to constrain docking and model selection, agreement between the model and these observations does not constitute independent structural validation.

Comparison with previously described VHL neosubstrates reveals both shared features and substrate-specific diversity. Although CDO1, GEMIN3 and caspase-2 show distinct structural degrons, in all three cases, target engagement appears to involve a composite surface that cooperate to form a glue-oriented interface with VHL^30,31^. The available data therefore suggest that VHL molecular glues can recruit proteins through multiple degron architectures rather than through a single conserved sequence or structural motif^30,31^. Moreover, as demonstrated in the reported CDO1 and GEMIN3 studies and in our analysis of caspase-2, cross-species domain swapping and targeted mutagenesis were critical for pinpointing degron motifs^30,31^, emphasizing the power of ortholog-guided approaches to define glue-dependent interaction surfaces.

An unexpected finding in our VHL-TurboID experiments was the robust enrichment of HIF-1α across dCASP2 compounds (**Fig. 1C-D**, **Fig. 3G-H**) despite minimal VHL-HIF-1α ternary-complex formation in the NanoBiT assay (**Supplementary Fig. S6**). Because these VHL-directed ligands mimic the hydroxyproline epitope within the HIF-1α oxygen-dependent degradation (ODD) motif, they are predicted to compete with, rather than promote, the native VHL-HIF-1α interface. Notably, both assays were performed under UPS inhibition to prevent protein turnover, indicating that the divergence reflects assay physics rather than degradation. We therefore propose two non-exclusive hypotheses that may account for this: (1) MGD-driven caspase-2-VHL assemblies re-orient VHL and reposition the TurboID enzyme so that it “sees” HIF-1α more frequently within its ∼10nm labeling radius, yielding increased time-integrated biotinylation rather than stabilizing VHL-HIF-1α interaction; (2) ligand occupancy of the VHL hydroxyproline pocket reduces HIF-1α dwell time at the native site (competition) in VHL, increasing the encounter frequency between VHL and HIF-1α, which in turn results in catalytic, time-integrating labeling by TurboID whereas NanoBiT requires a properly oriented, sustained interface.

The identification of selective caspase-2 degraders is particularly significant for the apoptosis and stress response fields. Although caspase-2 was one of the first caspases discovered, its biology has remained poorly understood due to the absence of selective pharmacological tools. Available caspase inhibitors lack specificity, and much of caspase-2 research has relied on genetic ablation models, complicating interpretation due to compensatory effects. Our degraders thus represent the first selective chemical probes for caspase-2, enabling temporal and dose-dependent control over its abundance. These tool degraders will allow for mechanistic studies of caspase-2 function in apoptosis, DNA damage response, and metabolic regulation, and may present therapeutic opportunities in neurodegenerative disease and other disorders where caspase-2 plays a role.

More broadly, our work demonstrates the potential of expanding molecular glue discovery beyond CRBN to other E3 ligases such as VHL. By demonstrating that VHL can be chemically reprogrammed to degrade a non-canonical substrate like caspase-2, we provide proof-of-principle that the neosubstrate landscape of VHL is broader than previously appreciated. These findings not only establish caspase-2 as a tractable target for VHL-based MGDs but also suggest the feasibility of discovering novel degrons and neosubstrates through unbiased proteomic approaches coupled with rational chemical optimization.

Together, our study establishes a foundation for exploring caspase-2 degraders as potential therapeutic agents. Caspase-2 has been implicated both as a tumor suppressor in certain malignancies and as a mediator of cellular stress responses. Accordingly, its selective modulation may offer therapeutic opportunities in oncology and degenerative diseases. While our current compounds primarily function as chemical probes, further medicinal chemistry optimization could improve their potency, selectivity, and pharmacokinetic properties, thereby facilitating translational development. The identification of caspase-2 as a novel neosubstrate for VHL expands the E3 ligase target landscape and provides a framework for the rational design of next-generation molecular glue degraders with therapeutic potential.

## Supplementary Figures

**Supplementary Figure S1.**
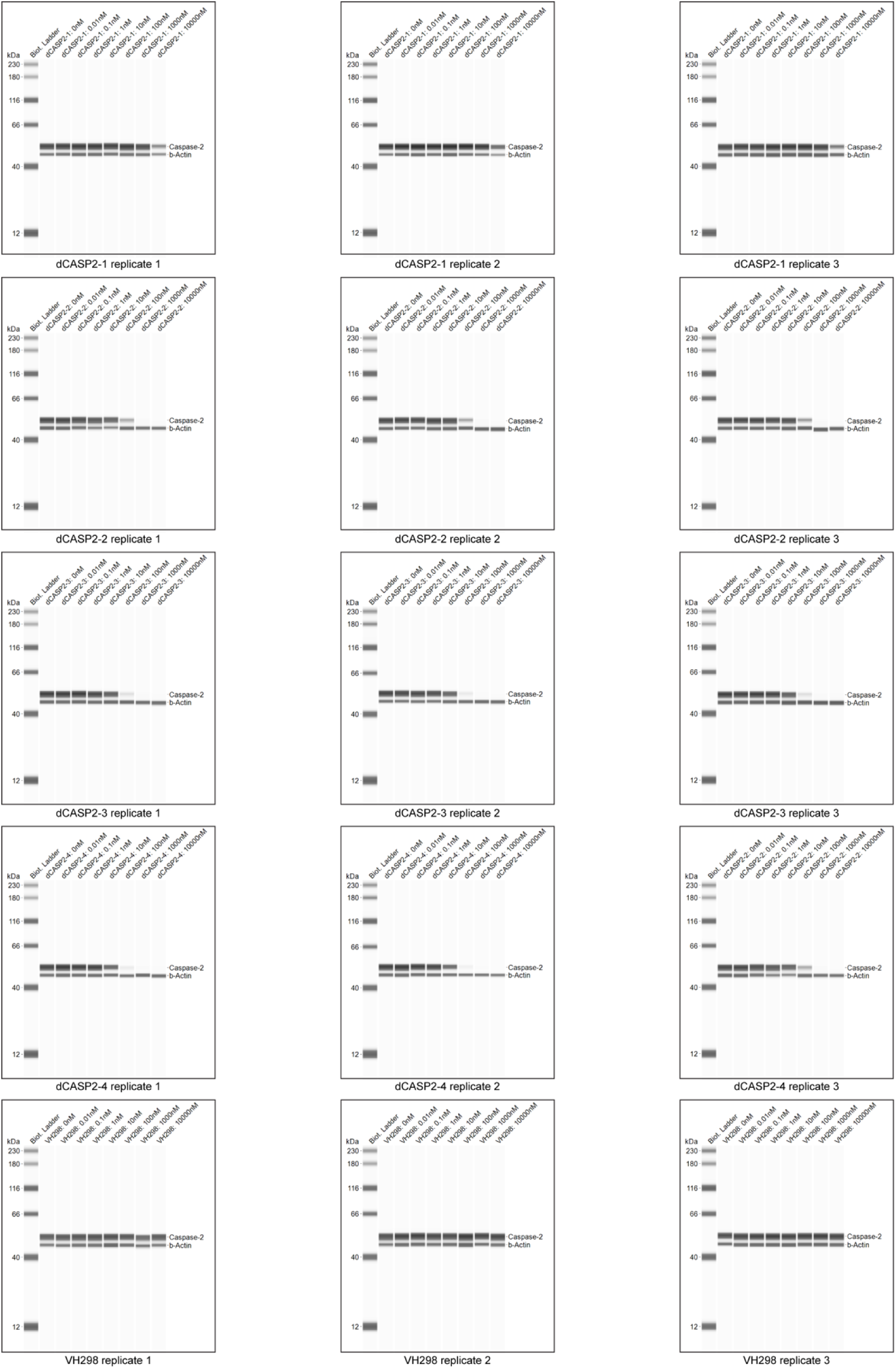
JESS Image for quantification of caspase-2 levels. Jurkat cell lysates were collected after 24-hour treatment with increasing concentrations of caspase-2 molecular glue degraders and negative control VH298. JESS quantification was performed in three biological replicates. β-actin was used as a loading control.

**Supplementary Figure S2.**
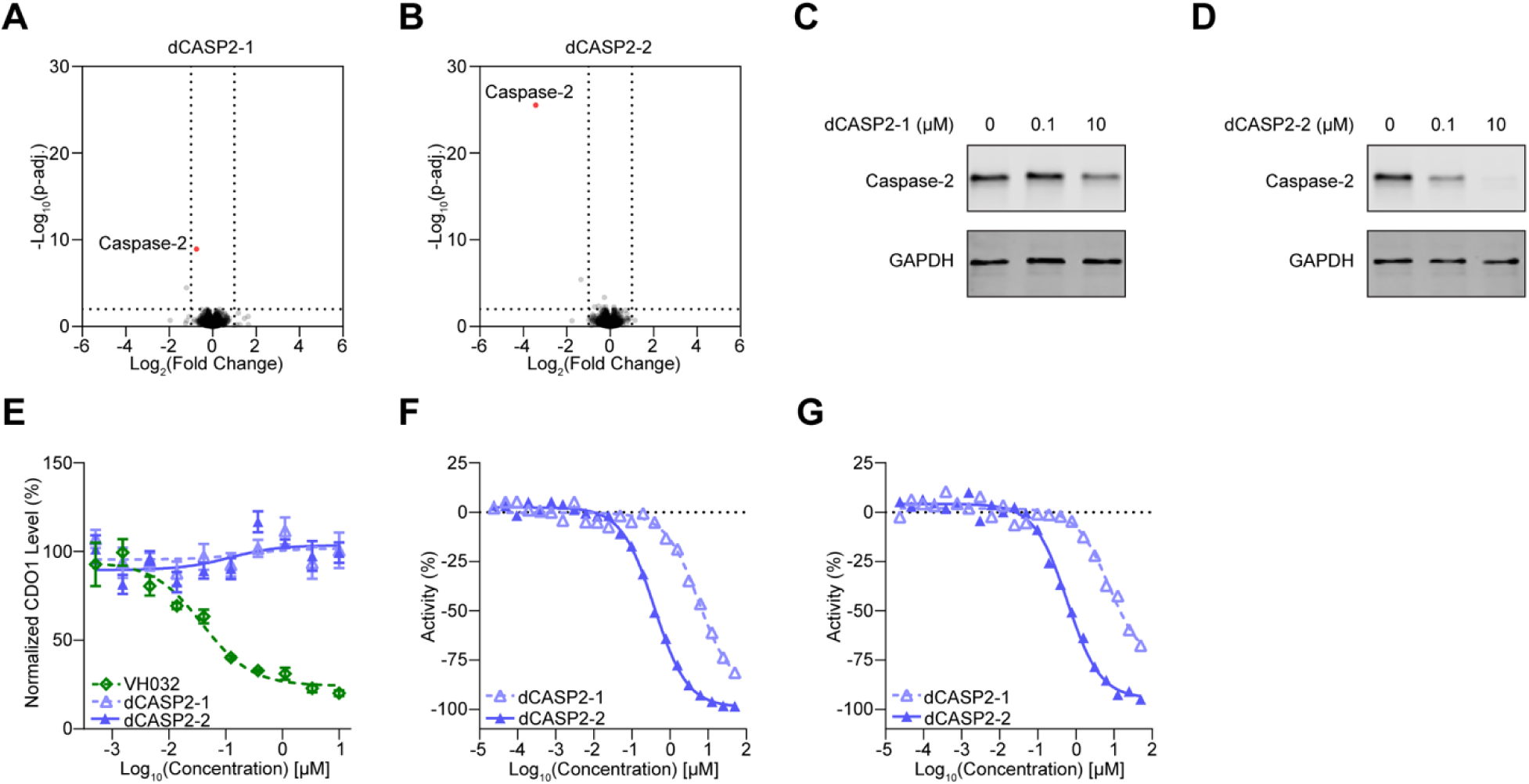
Rapid and selective caspase-2 degradation by dCASP2-1 and dCASP2-2. **A-B.** dCASP2-1 and dCASP2-2 promote caspase-2 downregulation. Wild-type Jurkat cells were treated for 6 hours with dCASP2-1 (10 μM, **A**, n=3), dCASP2-2 (10 μM, **B**, n=3), or DMSO (0.1%, n=3), followed by global proteomics analysis. Dashed lines indicate significance and enrichment cutoffs of adjusted p-value < 0.01 and log_2_ fold-change (relative to DMSO) < –1. Caspase-2 is highlighted in red. **C-D.** dCASP2-1 and dCASP2-2 rapidly degrade caspase-2. Jurkat cells were treated for 6 hours with increasing concentrations of dCASP2-1 (**C**) or dCASP2-2 (**D**). Endogenous caspase-2 was analyzed by Western blot, with GAPDH probed as a loading control. **E.** dCASP2-1 and dCASP2-2 are inactive in a CDO1 degradation assay. HiBiT assays were used to measure CDO1 levels after 24-hour treatment with dCASP2-1 or dCASP2-2. VH032 was included as a positive control (n=3). **F-G.** dCASP2-1 and dCASP2-2 bind to VHL. VBC NanoBRET VHL target engagement assays were performed in permeabilized (**F**) and live cells (**G**). Decreased % activity indicates increased VHL binding (n=1).

**Supplementary Figure S3.**
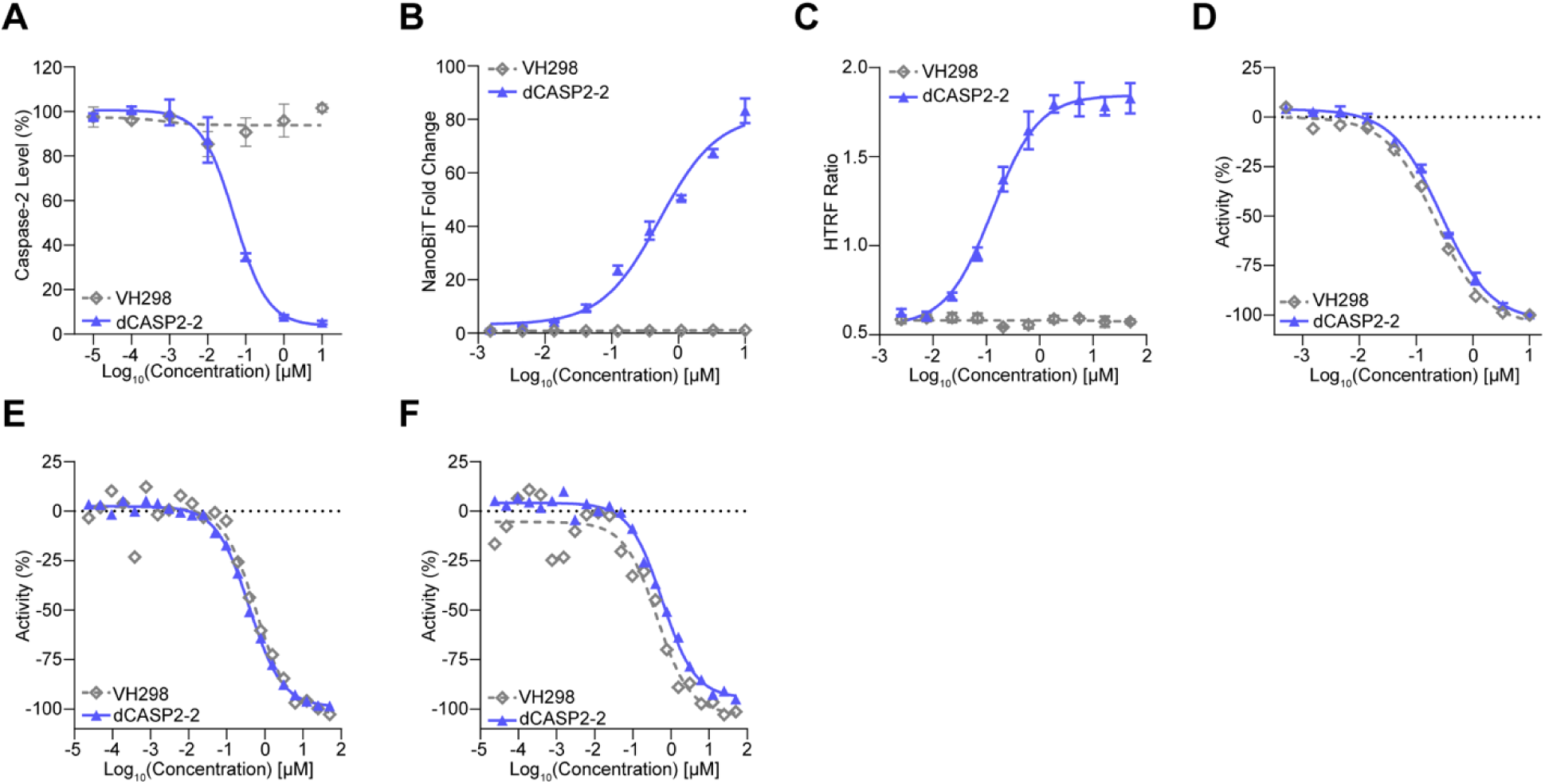
VH298 is not a caspase-2 degrader (dCASP2-2 was used as positive control). **A.** VH298 does not degrade caspase-2 in a dose-dependent manner. Caspase-2 levels were quantified by JESS following 24-hour VH298 treatment at increasing concentrations in Jurkat cells (n=3). **B.** VH298 does not induce caspase-2/MGD/VHL ternary complex formation in cells. HEK293T cells were transiently co-transfected with VHL-LgBiT and SmBiT-caspase-2 vectors, then co-treated with MLN4924 and caspase-2 degraders. Increased luminescence reflects compound-induced ternary complex formation (n=3). **C.** VH298 does not induce *in vitro* caspase-2/MGD/VHL ternary complex formation. HTRF assays were performed with purified caspase-2 and VBC complex in the presence of VH298. Higher HTRF signal indicates ternary complex formation *in vitro* (n=3). **D-F.** VH298 binds to VHL. VBC AlphaScreen competition assays were used to determine binary binding of VH298 to VHL *in vitro* (**D**). VBC NanoBRET VHL target engagement assays were also performed in permeabilized (**E**) and live cells (**F**). In all assays, decreased % activity indicates stronger VHL engagement (n=1).

**Supplementary Figure S4.**
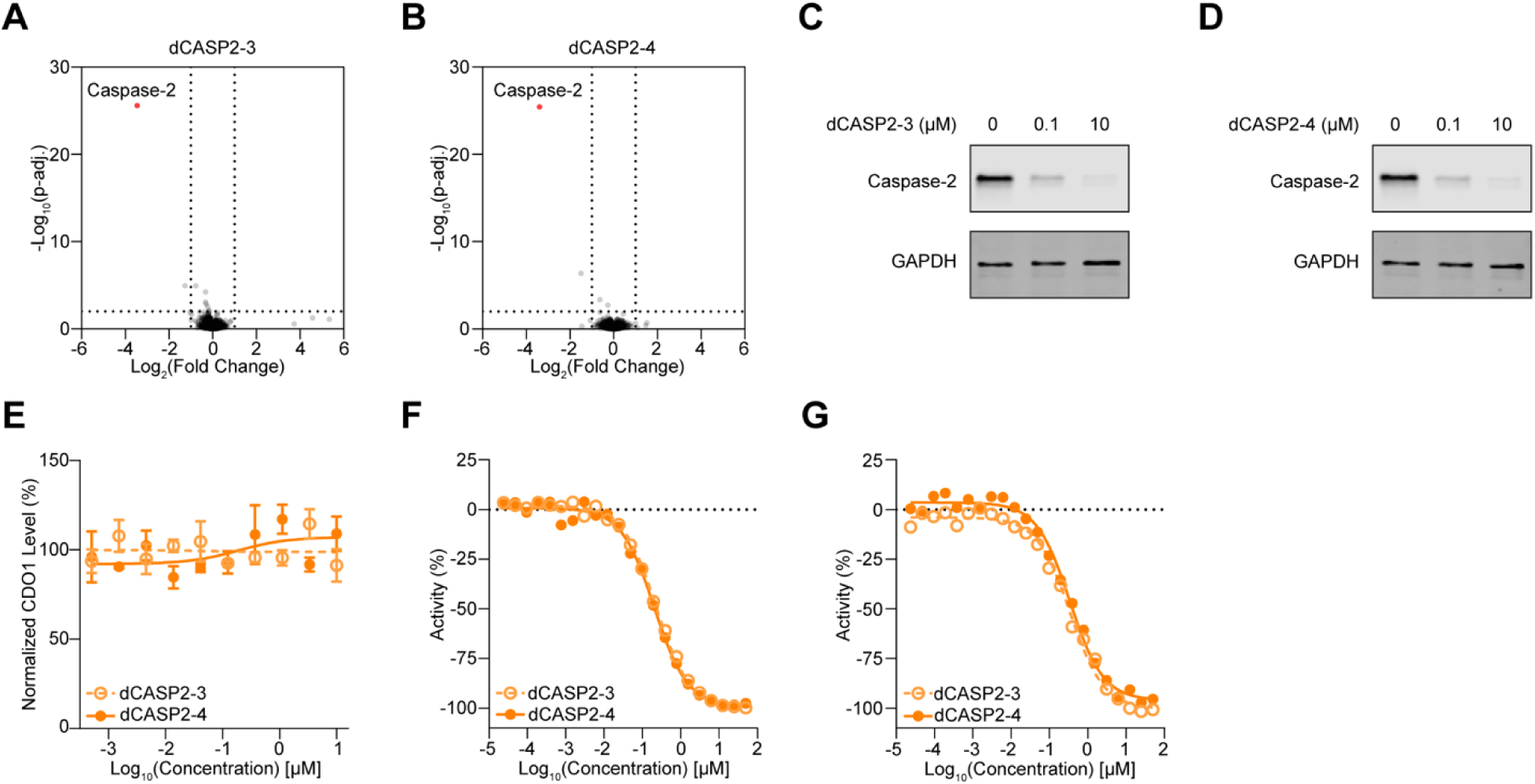
Rapid and selective caspase-2 degradation by dCASP2-3 and dCASP2-4. **A-B.** dCASP2-3 and dCASP2-4 promote caspase-2 downregulation. Wild-type Jurkat cells were treated for 6 hours with dCASP2-3 (10 μM, **A**, n=3), dCASP2-4 (10 μM, **B**, n=3), or DMSO (0.1%, n=3), followed by global proteomics analysis. Dashed lines indicate significance and enrichment cutoffs of adjusted p-value < 0.01 and log_2_ fold-change (relative to DMSO) < –1. Caspase-2 is highlighted in red. **C-D.** dCASP2-3 and dCASP2-4 rapidly degrade caspase-2. Jurkat cells were treated for 6 hours with increasing concentrations of dCASP2-3 (**C**) or dCASP2-4 (**D**). Endogenous caspase-2 was analyzed by Western blot, with GAPDH probed as a loading control. **E.** dCASP2-3 and dCASP2-4 are inactive in a CDO1 degradation assay. HiBiT assays were performed to measure CDO1 levels after 24-hour treatment with dCASP2-3 or dCASP2-4 (n=3). **F-G.** dCASP2-3 and dCASP2-4 bind to VHL. VBC NanoBRET VHL target engagement assays were performed in permeabilized (**F**) and live cells (**G**). Decreased % activity indicates stronger VHL binding (n=1).

**Supplementary Figure S5.**
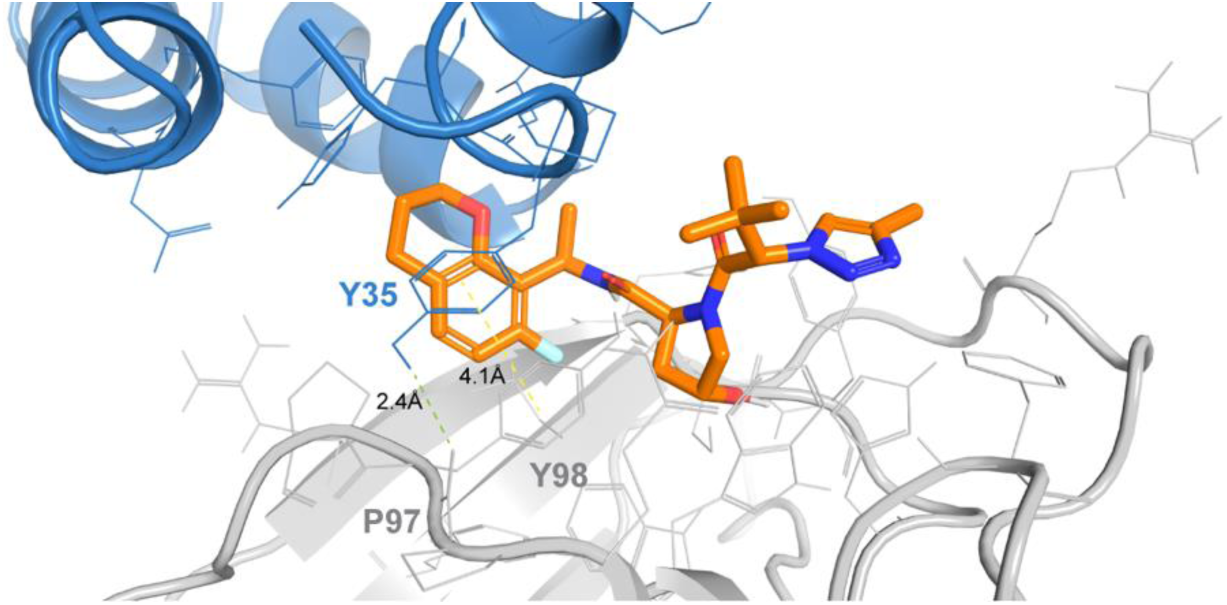
Computational 3D modeling predicts caspase-2 (H35Y)/dCASP2-3/VHL ternary complex. Selected computational model of ternary complex of caspase-2 carrying the H35Y substitution, dCASP2-3, and VHL. The mutant caspase-2 CARD domain is shown in blue, VHL in grey, and dCASP2-3 in orange. Key interactions between mutant caspase-2 Y35 and VHL (Y98, P97) are highlighted: green line represents hydrogen bond, and yellow line shows π–π aromatic stacking.

**Supplementary Figure S6.**
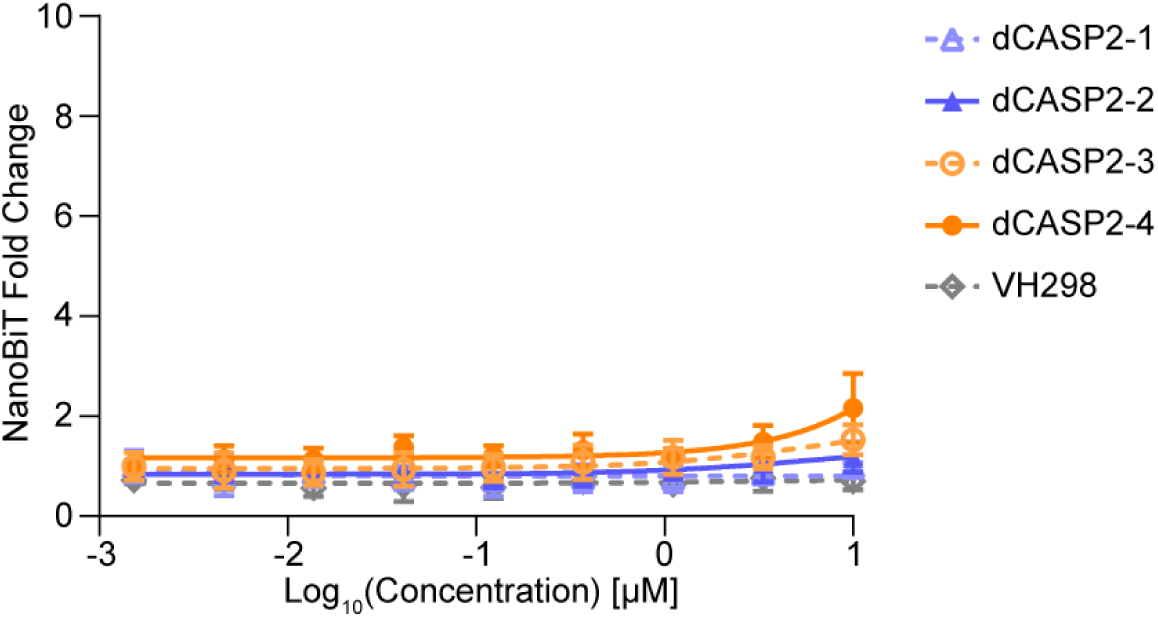
Cellular TCF assay for HIF-1α. NanoBiT assays were performed to measure compound-induced ternary complex formation between HIF1α and VHL.

## Methods

### Cell culture

Jurkat cells were maintained in RPMI1640 (Gibco #11875119) supplemented with 10% FBS (Gibco #16000044) and Penicillin-Streptomycin (Gibco # 15140122). HEK293T and RD cells were maintained in DMEM (Gibco #11965092) supplemented with 10% FBS (Gibco #16000044) and Penicillin-Streptomycin-Glutamine (Gibco #10378016), TrypLE Express dissociation reagent (Gibco #12605010) was used for HEK293T and RD cell passaging. All cells were incubated at 37°C with 5% CO_2_.

### Jurkat VHL TurboID cell line generation

Jurkat cells were passaged 2-3 days before transfection. For one-step VHL knockout-rescue, cells were electroporated with an sgVHL-1 CRISPR/Cas9 plasmid (to disrupt endogenous VHL, gRNA sequence: *CCGTCGAAGTTGAGCCATAC*) together with a PiggyBac donor encoding C-terminal TurboID-tagged VHL under a CAG promoter and a puromycin resistance marker and HyPBase. Neon™ NxT (10 µL tips, Invitrogen #N1096) was used for electroporation, 0.5 × 10⁶ cells in 10 µL Buffer R were used in each reaction with the settings of 1325 V, 10 ms, 3 pulses, the cells then were immediately transferred to 1 mL antibiotic-free medium per well in a well of a 12-well plate. After 24-hour incubation, puromycin 2 µg/mL (Gibco #A11138) was applied and refreshed every 2-3 days before single clones were isolated.

### TurboID proximity labeling

For TurboID proximity labeling cell treatment, Turbo-VHL expressing Jurkat cells were maintained in RPMI1640 (Gibco #11875119) supplemented with 10% FBS (Gibco #16000044) and Penicillin-Streptomycin (Gibco # 15140122). Cells were washed once with phosphate-buffered saline (DPBS; Gibco #14190136) and resuspended in Biotin-free medium (Biotin-Free RPMI1640+10% dialyzed FBS). Cells were diluted to 0.3 M/mL density and plated onto round-bottom 96-well plates (30,000 cells per well, Corning #3799). After two-day incubation at 37°C with 5% CO_2_, cells were treated with a mixture of Biotin (50 µM final concentration), Bortezomib (2 µM final concentration), and compound (10 µM final concentration) or DMSO control followed by 6-hour incubation at 37°C with 5% CO_2_. After incubation, cells were washed four times with DPBS (Gibco #14190136) using a BioTek® 405 Microplate Washer, and the cell pellets were stored at -80°C immediately.

For streptavidin enrichment and LC-MS sample preparation. Cell pellets were lysed in 150 µL of RIPA buffer (Thermo Scientific #89901) supplemented with 25 units of BenzoNuclease and a 1x Protease Inhibitor Cocktail (Thermo Scientific EDTA free, A32965). The cell lysates were subjected to sonication at 50 Hz for 20 minutes using a PIXUL sonicator (Active Motif) to ensure efficient cell disruption. For enrichment, 15 µL of high-capacity streptavidin magnetic beads (Thermo Scientific #88817) were added to the lysates, and the mixture was incubated to allow binding. Following incubation, the beads were sequentially washed six times with various buffers to remove non-specifically bound proteins: twice with RIPA buffer, once with 8 M Urea, once with 5 M NaCl, and twice with 50 mM EPPS buffer (stock: Thermo Scientific Chemicals #J61476.AK). Cleaned up beads were reduced and alkylated by adding Bond-Breaker TCEP (Thermo Scientific #77720) to a final concentration of 100 mM and Iodoacetamide (Thermo Scientific #A39271) to a final concentration of 100 mM and incubating at room temperature for 30 min. On-bead digestion was carried out by adding 0.5 µg of a Trypsin/Lys-C mixture (Promega #V5072) in 50 µL of 50 mM EPPS buffer per sample and incubating overnight at 37°C. Subsequently, 40% of each digested sample was loaded onto an Evotip (Evosep #EV2011) for downstream analysis.

For TurboID LC-MS data acquisition and analysis. Samples were analyzed on an EvoSep TIMSTOF-HT LC-MS system. Peptide separations were performed using the Evosep One 100SPD method (i.e. throughput of 100 samples per day) on an 8-cm PepSep C18 analytical column (Bruker #1893470). The TIMSTOF-HT operated in diaPASEF mode, with ion mobility separation spanning an inverse reduced mobility (1/K₀) range of 0.7-1.3 V·s/cm², mass range from 100-1700 m/z. Both the ramp time and accumulation time were set to 65 ms, and 36 scan windows were acquired per cycle, yielding a total cycle time of 0.85 s. diaPASEF data were searched against an in-house built Jurkat cell specific human spectral library using DIA-NN (v2.0.1 Enterprise), Mass accuracy and MS1 accuracy both were set to 15ppm, with match between runs enabled, normalization scheme was set to RT dependent. Protein group quant results were applied to statistical modeling and hypothesis testing using the limma package.

### Global proteomics

For global proteomics cell treatment, Jurkat wild type cells were maintained in in RPMI1640 (Gibco #11875119) supplemented with 10% FBS (Gibco #16000044) and Penicillin-Streptomycin (Gibco # 15140122). Cells were washed once with phosphate-buffered saline (DPBS; Gibco #14190136) and resuspended in the same culture media. Cells were diluted to 0.5 million/mL counting and plated onto round-bottom 96-well plates (50,000 cells per well, Corning #3799). After overnight incubation at 37°C with 5% CO2, cells were treated with compound (10 µM final concentration) or DMSO control followed by 6- or 24-hour incubation at 37°C with 5% CO2. After incubation, cells were washed four times with DPBS (Gibco #14190136) using a BioTek® 405 Microplate Washer, and the cell pellets were stored at -80°C immediately.

For global proteomics sample preparation, cell pellets were lysed in 100 µL of lysis buffer (2% SDS, 100 mM Tris-HCl) and sonicated using an Abcam PIXUL sonicator (50Hz, 20 min). The protein concentration of the lysates was determined by the BCA method (Thermo Scientific #23225). For each sample, 30µg of protein was reduced and alkylated by adding Bond-Breaker TCEP to a final concentration of 100 mM and chloroacetamide to a final concentration of 500 mM and incubating at 56 °C for 45 min. The samples were then subjected to the SP3 protocol for clean-up. Briefly, 30 µg of protein was bound to >300 µg of beads in 80% (v/v) ethanol for 15 min and then washed three times with 80% ethanol and once with acetonitrile. The protein-bound beads were digested with 1.2 µg of Trypsin/Lys-C mix in 50 mM EPPS for 18 h at 37 °C. The beads were removed, and the 800ng of digested peptides were loaded and cleaned up onto Evotip for downstream analysis.

For global proteomics data acquisition and analysis, samples were analyzed on an EvoSep Orbitrap Astral LC-MS system. Peptide separations were performed using the Evosep One 100SPD method (i.e. throughput of 100 samples per day) on an 8-cm Aurora Rapid 8 x 150 XT C18 (IonOptics, #AUR3-80150C18XT) analytical column. On the Orbitrap Astral mass spectrometer, data were acquired in DIA mode. MS1 full scans were collected in the Orbitrap at a resolution of 240,000 (at m/z 200) over a scan range of 380-980 m/z, with a normalized AGC target of 500% and a maximum injection time of 5 ms. MS2 spectra were acquired in the Astral mass analyzer using a precursor mass range of 380-980 m/z, with a 2 m/z isolation window and no window overlap. Fragmentation was performed using higher-energy collisional dissociation (HCD) with a normalized collision energy of 25%. DIA data were searched against an in-house built Jurkat cell specific human spectral library using DIA-NN (v2.0.1 Enterprise), Mass accuracy was set to 10ppm and MS1 accuracy was set to 4ppm, with match between runs enabled, normalization scheme was set to RT dependent. Protein group quant results were applied statistical modeling and hypothesis testing using the limma package with Benjamini-Hoechberg muti-test correction.

### Measurement of protein level change by western blotting

Cells treated with compound and control were harvested and washed once with ice-cold DPBS (Gibco #14190136). Cell pellets were lysed with radioimmunoprecipitation assay lysis and extraction buffer (Thermo Scientific #89901) supplemented with protease and phosphatase inhibitor cocktail (Thermo Scientific #78440). Protein lysates were incubated on ice with gentle shaking for 15 min before being centrifuged at 4°C/15,000 rpm for 15 min. Supernatants were transferred into new centrifuge tubes, and an equal volume of 2-mercaptoethanol (Bio-Rad #1610710) supplemented 2× Laemmli sample buffer (Bio-Rad #1610737) was added to the protein lysate. Protein samples were heated at 99°C before being stored at −20°C in the freezer or for Western blotting. Equal amounts of protein samples were run on precast 4 to 15% tris-glycine Mini-or Midi-PROTEAN TGX gels (Bio-Rad #4561086, 5671085). Resolved proteins were transferred onto nitrocellulose membranes using Bio-Rad Turbo transfer system (Bio-Rad #1704271, 1704271). The membranes were blocked with Intercept (tris-buffered saline) blocking buffer (LI-COR #927-60001) for 1 hour at room temperature. The membranes were then probed with primary antibodies (Abcam #ab179520, Cell Signaling #97166) at an optimal concentration [dilution factor of 1:1000 for caspase-2 and 1:10,000 for glyceraldehyde-3-phosphate dehydrogenase (GAPDH)] in the antibody diluent (LI-COR #927-65001) overnight at 4°C. The membranes were washed with TBST three times (10 min each time on a shaker) and incubated with the Anti-rabbit IgG (H+L, DyLight 800 Conjugate) and Anti-mouse IgG (H+L, DyLight 680 Conjugate) secondary antibodies (Cell Signaling #5470, 5151) at room temperature for 2 hours. Excessive antibodies were washed with TBST, and the membranes were exposed by LI-COR Odyssey imaging system. Immunoblotting data were analyzed with Image Studio software (Li-Cor, Version 5.5).

### Measurement of protein level change by JESS

Cell lysates were analyzed with the Jess Automated Western Blot System (ProteinSimple #004-650) according to the manufacturer’s instructions using 12–230 kDa Separation Modules with 25 Capillary Cartridges (ProteinSimple #SM-W004) and Anti-Rabbit Detection Module (ProteinSimple #DM-001). Prepared lysates at 2X working concentration (1 mg/mL) were diluted 1:1 with 0.1X Sample Buffer and combined with 5X Fluorescent Master Mix in a 1:5 ratio to make final lysate concentrations of 0.4 mg/mL. 1.2 μg protein per sample, antibody diluent, primary antibodies in antibody diluent, HRP-conjugated secondary antibodies and chemiluminescent substrate were pipetted into the 12-230 kDa Pre-Filled Microplate. Instrument default settings were used: stacking and separation at 375 V for 25 min; blocking reagent for 5 min, primary and secondary antibody both for 30 min; Luminol/peroxide chemiluminescence detection for ∼15 min. The resulting electropherograms were inspected to check whether automatic peak detection required any manual correction using Compass for SW (ProteinSimple, Version 6) software. The primary antibodies used were anti-caspase-2 and anti- β-actin (Abcam #ab179520 and Cell Signaling #5125). Secondary antibodies used were Anti-Rabbit-HRP (Cell Signaling #7074, 1:50) for caspase-2 detection. Anti-Rabbit Detection Module (ProteinSimple #DM-001) also contained the following reagents used for all gel capillary electrophoresis set up: Antibody Diluent 2, Luminol-S chemiluminescence detection, and Peroxide chemiluminescence detection. Acquired data were subsequently analyzed with Compass for SW (ProteinSimple, Version 6) software.

### HTRF ternary complex formation assay

To measure the compound-induced interaction between VHL and caspase-2, we performed a homogenous time-resolved fluorescence (HTRF) assay. 30 nM purified recombinant biotinylated VBC complex (Viva Biotech) was mixed with 40 nM His-SUMO-caspase-2 (CUSABIO #CSB-EP004547HU), 1X streptavidin-terbium cryptate (Revvity #610SATLB), and 1X anti-6His-d2 (Revvity #61HISDLB) in assay buffer (PBS (Gibco #14190-136) with 0.1% BSA (Sigma #BP9704), 0.01% Tween-20 (ThermoFisher #28352), and 1 mM DTT (Thermo Scientific #20290). 20 μL reactions were dispensed into a Proxiplate 384-shallow well (Revvity #6008289). 100 nL of each compound concentration was added to the reactions by acoustic dispensing of stocks in 100% DMSO. The reactions were incubated at room temperature overnight, and then the plate was read on an EnVision Xcite. The resulting HTRF ratio (665/620) was plotted in dose-response format to evaluate ternary complex formation.

### HIF-1α-VBC AlphaScreen competition binding assay

For AlphaScreen, the recombinant VHL-Elongin B/C (VBC) complex was purified by immobilized metal-affinity chromatography followed by size-exclusion chromatography. Constructs encoding VHL (54-213), ElonginB (17-112), ElonginC (1-104) were used to express the VBC complex. Cells were resuspended in lysis buffer (20 mM HEPES, 250 mM NaCl, and 2 mM imidazole) and lysed by microfluidization. Following clarification by ultracentrifugation, the supernatant was incubated with Talon resin (Clontech, #635507) for 2 hours at 4°C, washed extensively, and the bound VBC complex was eluted with 250mM imidazole. Eluted protein was then purified using Superdex S-75 column (GE Healthcare) and stored in buffer (10mM HEPES 7.5, 150mM NaCl, 0.5mM TCEP) at - 80°C prior to AlphaScreen assay. HIF-1α-VBC AlphaScreen competition binding assay was performed in a 15 µL volume of assay buffer (50 mM Tris, pH 7.5 / 150 mM NaCl / 1 mM TCEP / 0.1% BSA) in a Corning 384-well low flange white polystyrene NBS microplate (Corning #3574). In concentration-response experiments with tested compounds, 0.5 µL of 10 concentrations from 3-fold serial dilutions in DMSO with a top concentration of 0.3 mM was pre-incubated with 7 µL of recombinant His-tagged VBC at 8.58 nM for 10 minutes at room temperature. Then, 7.5 µL of biotinylated HIF-1a Ciulli peptide at 6 nM was added and incubated for 30 minutes at room temperature. Finally, 15 µL of 2-bead combo (nickel chelate donor beads (Revvity #AS101) and streptavidin acceptor beads (Revvity #AL125) at 40 µg/mL in 50 mM Tris, pH 7.5 / 150 mM NaCl / 0.2% Tween-20 / 0.2% BSA) was combined with the above binding mixture and incubated for 60 minutes in dark. The luminescence signal of each well was measured using an EnVision multilabel plate reader (Waltham, MA) with a default EnVision AlphaScreen assay protocol. The final concentrations of recombinant His-tagged VBC, biotinylated HIF-1a Ciulli peptide, nickel chelate donor beads and streptavidin acceptor beads are 4 nM, 3 nM, 20 µg/mL and 20 µg/mL, respectively in the assay.

### VHL permeabilized cell and live cell NanoBRET target engagement assays

HEK293T cells were maintained in DMEM (Gibco #11965092) supplemented with 10% FBS (Gibco #16000044) and Penicillin-Streptomycin-Glutamine (Gibco #10378016). For construct expression, 4 million cells were transfected with 9 µg Transfection Carrier DNA (Promega #E488A) and 1 µg VHL-NanoLuc Fusion Vector (Promega #N275A) using 30 µL FuGENE HD Transfection Reagent (Promega #E2312) according to the manufacturer’s protocol. After the transfection, cells were incubated at 37°C overnight. Test compounds were dispensed into 384-well ProxiPlates (Revvity #6008280) using an Echo650 acoustic dispenser (Beckman Coulter). Cells were then lifted and resuspended to a concentration of 500,000 cells/mL in Opti-MEM (Gibco #11058021) supplemented with 1% (w/v) FBS, 4% (w/v) Tracer Dilution Buffer (Promega #N219A), VHL NanoBRET Tracer (Promega #N292A; 0.25 µM VHL Tracer for permeabilized format or 0.5 µM VHL Tracer for live format), and 50 ug/mL digitonin (Sigma #D141; digitonin was only included in the permeabilized cell assay). Cells were dispensed (5000 cells in 10 µL per well) using a Multidrop Combi dispenser (Thermo Scientific). The cells were then centrifuged at 500 RPM for 1 minute and mixed at 350 RPM for 30 seconds on an orbital shaker. For the permeabilized cell assays, NanoBRET Nano-Glo Substrate (Promega #N157A) was immediately added to the cells following the manufacturer’s protocol using a Multidrop Combi dispenser (Thermo Scientific); the cells were then centrifuged at 500 RPM for 1 minute, mixed at 350 RPM for 30 seconds on an orbital shaker, and incubated at room temperature for 5 minutes. After incubation, NanoBRET donor and acceptor signals were measured using the EnVision plate reader (Revvity). For the live cell assays, plates containing cells and test compounds were further incubated at 37°C for 2 hours. After the incubation, the cells were equilibrated to room temperature for 15 minutes. NanoBRET Nano-Glo Substrate (Promega #N157A) was added to the cells following the manufacturer’s protocol using a Multidrop Combi dispenser (Thermo Scientific); the cells were then centrifuged at 500 RPM for 1 minute, mixed at 350 RPM for 30 seconds on an orbital shaker, and incubated at room temperature for 5 minutes. After incubation, NanoBRET donor and acceptor signals were measured using the EnVision plate reader (Revvity). The EnVision plate reader was equipped with a 460/80 bandpass filter for detecting donor signals and a 647/75 bandpass filter for detecting acceptor signals. BRET signals were measured within 1h after adding substrate solution with a 1.0 s integration time. The BRET ratio was defined as the ratio of acceptor signal over the donor signal.

### NanoBiT assay

cDNA for *VHL*, *HIF1A*, and *CASP2* were synthesized, subcloned using SgfI/PmeI restriction sites into NanoBiT PPI Flexi vectors (Promega #N2015). HEK293T cells were plated at 10k cells per well in 96-well plates in FluoroBrite™ DMEM (Gibco #A1896701) supplemented with 5% FBS and GlutaMAX™ supplement (Gibco #35050061). After cells adhered to the plates, transfection mixtures were prepared in bulk and each well was individually transfected using 0.3 μL Fugene HD (Promega #E2311), 50 ng LgBiT-tagged VHL, and 50 ng SmBiT-tagged target in 5 μL OptiMEM. After 24-hour transfection, a 30-minute pre-treatment with 1 μM MLN4924 was used to inhibit protein degradation, followed by treatment of increasing concentrations of compounds. After 4-hour incubation, plates were brought to room temperature, and 25 μL of freshly prepared Nano-Glo Live Cell Reagent (Promega #N2012) was added to each well. Luminescence was measured immediately after mixing using an EnVision plate reader.

### CDO1 degradation with HiBiT assay

For the CDO1 degradation assay, HiBiT-CDO1 RD cells were dispensed in 384-well format with 2,000 cells per well and cultured overnight. Compounds were dosed by acoustic dispensing (Echo 555) of 10mM DMSO stocks, and HiBiT-tagged proteins were detected by addition of Nano-Glo HiBiT Lytic Reagent (Promega #N3040) according to the manufacturer specifications and measuring luminescence on a multimode plate reader (EnVision Xcite).

### Caspase2 degron mapping with NanoBRET assay

Constructs were generated with Flexi vectors using SgfI/PmeI restriction sites (Promega #G2821) to express NanoLuc-tagged caspase-2 variants (donor) and Halo-tagged VHL (acceptor). HEK293T cells transiently transfected with donor and acceptor fusion constructs using FuGENE HD according to the manufacturer’s protocol (Promega, NanoBRET™ Protein:Protein Interaction System Technical Manual). After ∼20 h expression, cells were replated into white 96-well plates at a density of 0.2M cells/ml and incubated with 100 nM HaloTag^®^ NanoBRET™ 618 Ligand or DMSO vehicle control. Compounds were added at 10× stock to yield final assay concentrations, and cells were incubated for 6 hours at 37°C before measurement. NanoBRET™ Nano-Glo^®^ Substrate was added to each well, and luminescence was recorded using dual-filtered detection of a EnVision plate reader at 460 nm (donor) and 618 nm (acceptor). Corrected NanoBRET ratios (mBU) were calculated by subtracting no-ligand control values from ligand-treated samples.

### Computation modeling for binary and ternary complex

Structure preparation: The VHL-Elongin B/C (VBC) complex (PDB: 9BOL) was used as the receptor for VHL. dCASP2 analogs were built with explicit stereochemistry and cyclization states. The caspase-2 CARD domain was obtained from the AlphaFold Protein Structure Database. Protein and ligand structures were prepared using standard Schrödinger workflows prior to docking and molecular dynamics (MD) simulations. Apo VHL structure (1VCB) was used as a reference to measure the conformational change upon dCASP-2 binding.

Protein-ligand docking (VHL binaries): Docking of dCASP2 analogs into VBC was performed with Schrödinger Glide^38^, enforcing canonical VHL hydrogen-bond constraints. Top-ranked binding poses were advanced into explicit-solvent MD simulations. Persistence of key VHL contacts (Y98, H110, S111, H115) and ligand RMSD relative to the starting pose were monitored to assess binding-mode fidelity and conformational stability across the degrader series.

Ternary complex modeling: Binary VHL-dCASP2 models were docked against the caspase-2 CARD domain using ensemble protein-protein docking with PIPER^40^. Conformational ensembles for both VHL and caspase-2 were generated from MD trajectories to capture relevant flexibility. Based on degron-mapping results, attractive restraints were applied to CARD domain residues H33, P34, H35, and D100. From ∼1,000 initial docking solutions, 10-15 representative complexes were selected for refinement and subjected to extended MD (≥200 ns) to evaluate interface stability, hydrogen-bonding, and salt-bridge interactions.

Free-energy perturbation (FEP): Contribution of individual caspase-2 degron residues was further assessed using Schrödinger FEP+^41^. Mutations H33Q, D100S, and H35Y were simulated in the dCASP2-3 ternary model (P34R excluded due to convergence limitations). Predicted ΔΔG values were compared qualitatively with experimental mutational data to validate the modeled interaction network.

## Chemistry Supplement

### General Synthetic Scheme

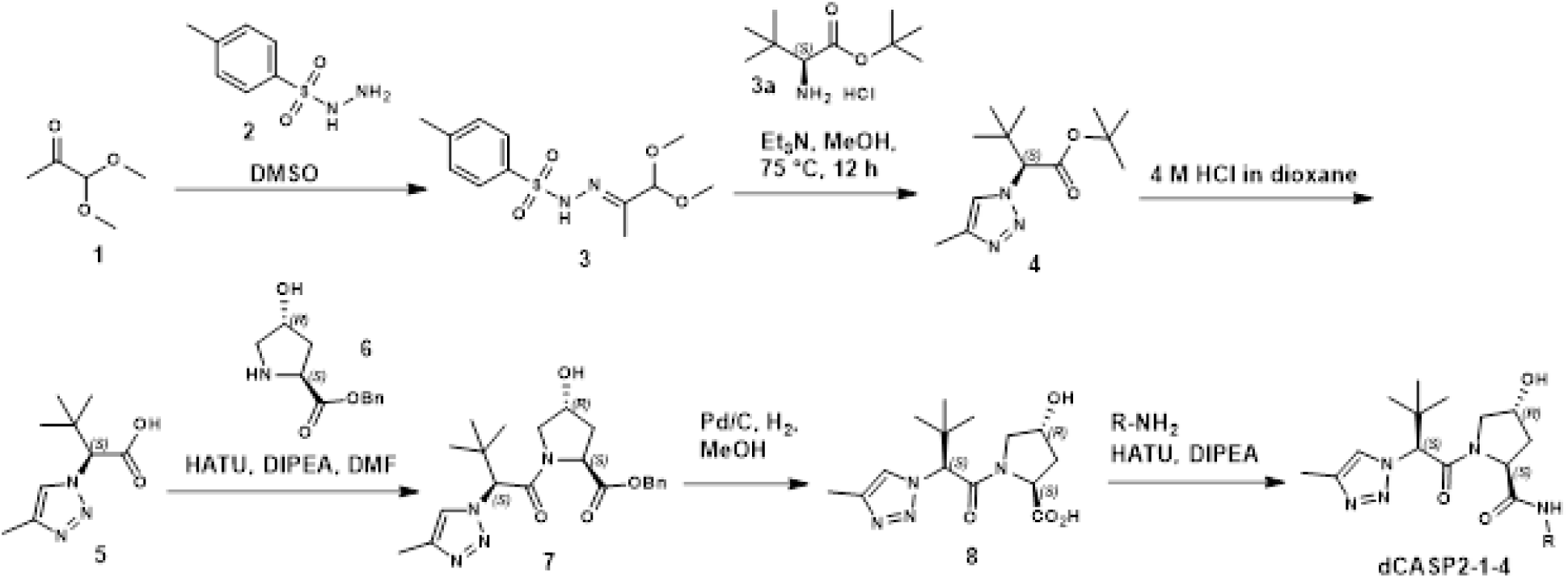

### Step 1: Synthesis of N’-(1,1-dimethoxypropan-2-ylidene)-4-methyl benzene sulfonohydrazide (3)

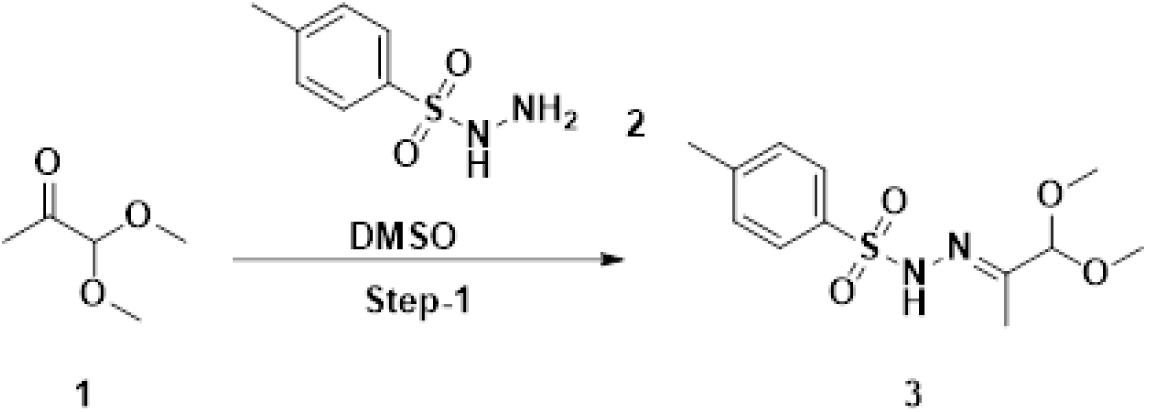

To a stirred solution of 4-methylbenzenesulfonohydrazide (**2**, 22 g, 118 mmol, 1.0 equiv) in dimethyl sulfoxide (100 mL) was added 1,1-dimethoxypropan-2-one (**1**, 14.44 mL, 119 mmol, 1.01 equiv) dropwise at 25 °C under nitrogen atmosphere and stirred for 1 h. The reaction mixture was diluted with ice cold water (500 mL) and the precipitated solid was filtered and washed with water (200 mL). The solid cake was dried under vacuum and triturated with hexane (300 mL) to give N’-(1,1-dimethoxypropan-2-ylidene)-4-methylbenzenesulfonohydrazide (**3**, 30 g, 94 mmol, 80% yield) as a white solid.

**LCMS (ESI, Positive ion) *m/z*:** 287.25 (M+H)^+^

**^1^H-NMR (400 MHz, DMSO- *d₆*):** δ 10.45 (s, 1H), 7.74 (d, *J* = 8.4 Hz, 2H), 7.39 (d, *J* = 8.0 Hz, 2H), 4.36 (s, 1H), 3.14 (s, 6H), 2.37 (s, 3H), 1.69 (s, 3H).

### Step-2: Synthesis of *tert*-butyl (S)-3,3-dimethyl-2-(4-methyl-1H-1,2,3-triazol-1-yl)butanoate

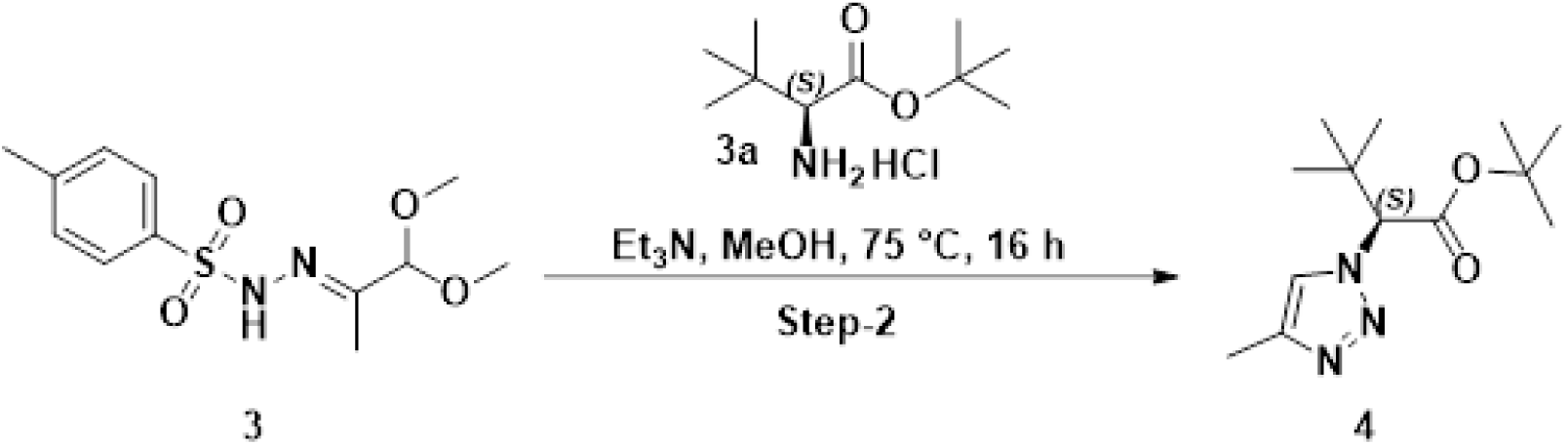

To a stirred solution of N’-(1,1-dimethoxypropan-2-ylidene)-4-methylbenzenesulfono-hydrazide (**3**, 20 g, 62.9 mmol, 1.0 equiv) in methanol (200 mL) were added *tert*-butyl (*S*)-2-amino-3,3-dimethylbutanoate hydrochloride (**3a**, 15.47 g, 69.1 mmol), 1.1 equiv) and triethyl amine (26.3 mL, 94 mmol, 3.0 equiv) at 25 °C under nitrogen atmosphere. The reaction mixture was stirred at 75 °C for 16 h. The reaction mixture was concentrated under reduced pressure. The crude compound was purified by column chromatography (silica: 100-200 mesh, elution: 0 to 20% ethyl acetate in hexane) to give (*tert*-butyl (S)-3,3-dimethyl-2-(4-methyl-1H-1,2,3-triazol-1-yl)butanoate (**4**, 9.1 g, 34.8 mmol, 55% yield) as an off white solid. Observed [a]: +15.48 (c= 0.48, MeOH).

**LCMS (ESI, Positive ion) *m/z*:** 254.36 (M+H)^+^

**^1^H-NMR (400 MHz, DMSO- *d₆*:** δ 7.95 (s, 1H), 5.10 (s, 1H), 2.25 (s, 3H), 1.44 (s, 9H), 0.97 (s, 9H).

**Chiral Purity:** 100 % ee

**Chiral Purity Method:**

Column/dimensions: Chiralcel OX-H (4.6 x 250) mm, 5μ, CO2%: 85%, Co-Solvent % :15% (0.5% DEA in MeOH), Total Flow :3 mL/min, Back Pressure :1500PSI, Temperature :30 °C

### Step-3: Synthesis of (*S*)-3,3-dimethyl-2-(4-methyl-1*H*-1,2,3-triazol-1-yl)butanoic acid

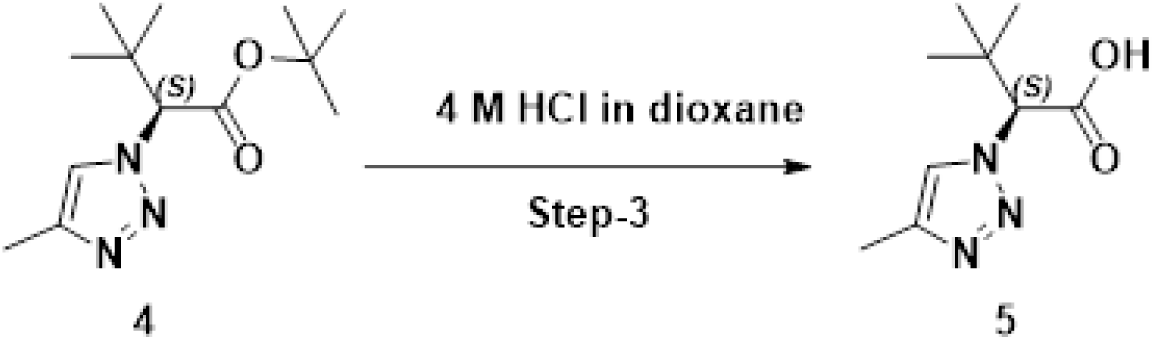

To a solution of *tert*-butyl (*S*)-3,3-dimethyl-2-(4-methyl-1H-1,2,3-triazol-1-yl)butanoate (**4**, 9.0g, 27.3 mmol, 1.0 equiv) in dichloromethane (20 mL) was added HCl (4M in dioxane, 90.17 mL, 361 mmol, 10.0 equiv) at 0 °C under nitrogen atmosphere and stirred at 55 °C for 6 h. The reaction mixture was concentrated under reduced pressure. The crude residue was triturated with diethyl ether (100 mL) and dried under vacuum to give (*S*)-3,3-dimethyl-2-(4-methyl-1H-1,2,3-triazol-1-yl)butanoic acid (**5**, 5.0 g, 15.54 mmol, 57% yield) as a gummy solid.

**LCMS (ESI, Positive ion) *m/z*:** 198.4 (M+H)^+^

**^1^H-NMR (400 MHz, DMSO-*d₆*:)** δ 7.95 (s, 1H), 5.13 (s, 1H), 2.25 (s, 3H), 0.98 (s, 9H).

**Chiral Purity:** 96.3 % ee

**Chiral Purity Method:**

Column/dimensions: Chiralcel OX-H (4.6 x 250) mm, 5μ, CO2%: 85%, Co-Solvent % :15% (0.5% DEA in MeOH), Total Flow :3 mL/min, Back Pressure :1500PSI, Temperature :30 °C

### Step-4: Synthesis of benzyl (2*S*,4*R*)-1-((S)-3,3-dimethyl-2-(4-methyl-1H-1,2,3-triazol-1-yl)butanoyl)-4-hydroxypyrrolidine-2-carboxylate

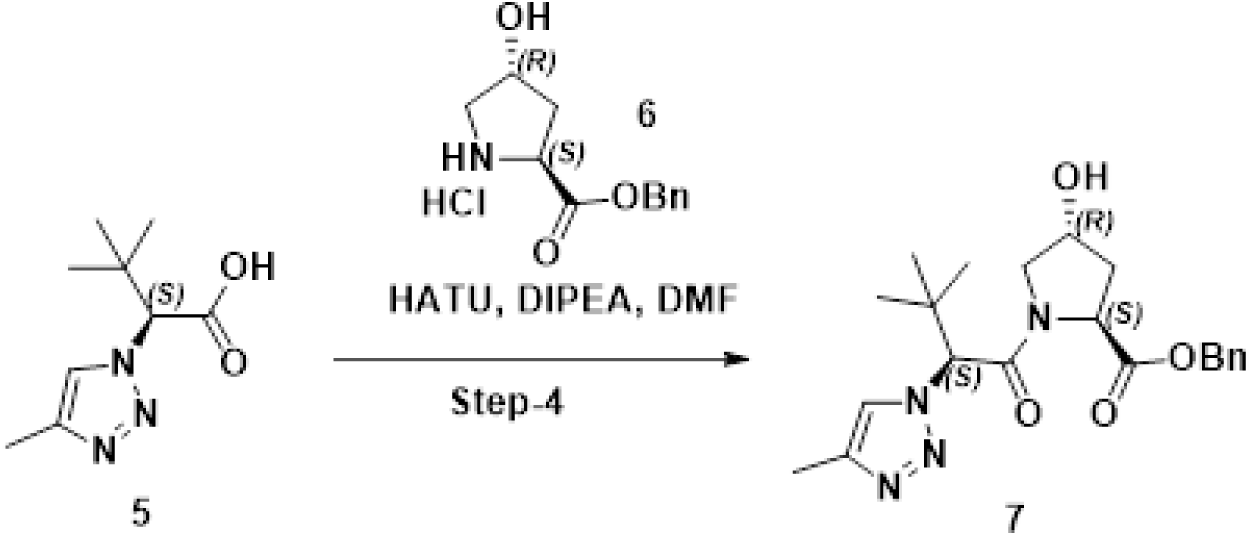

To a stirred solution of (*S*)-3,3-dimethyl-2-(4-methyl-1H-1,2,3-triazol-1-yl)butanoic acid (**5**, 3.0 g, 13.95 mmol, 1.0 equiv) in N,N-dimethylformamide (30 mL) was added DIPEA (14.62 mL, 84 mmol, 6.0 equiv) followed by HATU (7.96 g, 20.93 mmol, 1.5 equiv) and benzyl (2S,4R)-4-hydroxypyrrolidine-2-carboxylate hydrochloride (**6**, 3.60 g, 13.95 mmol, 1.0 equiv) at 0 °C under nitrogen atmosphere. The reaction mixture was stirred at 25 °C for 5 h. The reaction mixture was poured into ice cold water (150 mL). The precipitated brown solid was filtered and washed with water (20 mL), dried under vacuum and triturated with n-pentane (100 mL) to afford benzyl (2*S*,4*R*)-1-((*S*)-3,3-dimethyl-2-(4-methyl-1*H*-1,2,3-triazol-1-yl)butanoyl)-4-hydroxypyrrolidine-2-carboxylate (**7**, 5.21 g, 8.00 mmol, 57% yield) as an off white solid. Reported [a]: -35.5 (c = 1, MeOH). Observed [a]: -34.40 (c = 1, MeOH).

**LCMS (ESI, Positive ion) *m/z*:** 401.31 (M+H)^+^

**^1^H-NMR (400 MHz, DMSO-*d₆*):** δ 7.98 (s, 1H), 7.38 – 7.33 (s, 5H), 5.44 (s, 1H), 5.23 (d, *J* = 3.6 Hz, 1H), 5.22 (s, 2H), 4.35 – 4.42 (m, 1H), 3.78 (dd, *J* = 10.8, 4.0 Hz, 1H), 3.62 (d, *J* = 10.8 Hz, 1H), 2.23 (s, 3H), 2.12 – 2.19 (m, 2H), 1.91 – 1.99 (m,1H), 0.93 (s, 9H).

**Chiral Purity:** 81.1 % ee

**Chiral Purity Method:**

Column/dimensions: Chiralcel OX-H (4.6 x 250) mm,5μ% CO2 :70%, % Co solvent :30% (0.2% 7N Methanolic ammonia in ACN:MeOH)(1:1), Flow: 3mL/min, Back Pressure :1500 psi, Temperature :30°C

### Step-5: Synthesis of (2*S*,4*R*)-1-((*S*)-3,3-dimethyl-2-(4-methyl-1*H*-1,2,3-triazol-1-yl)butanoyl)-4-hydroxypyrrolidine-2-carboxylic acid

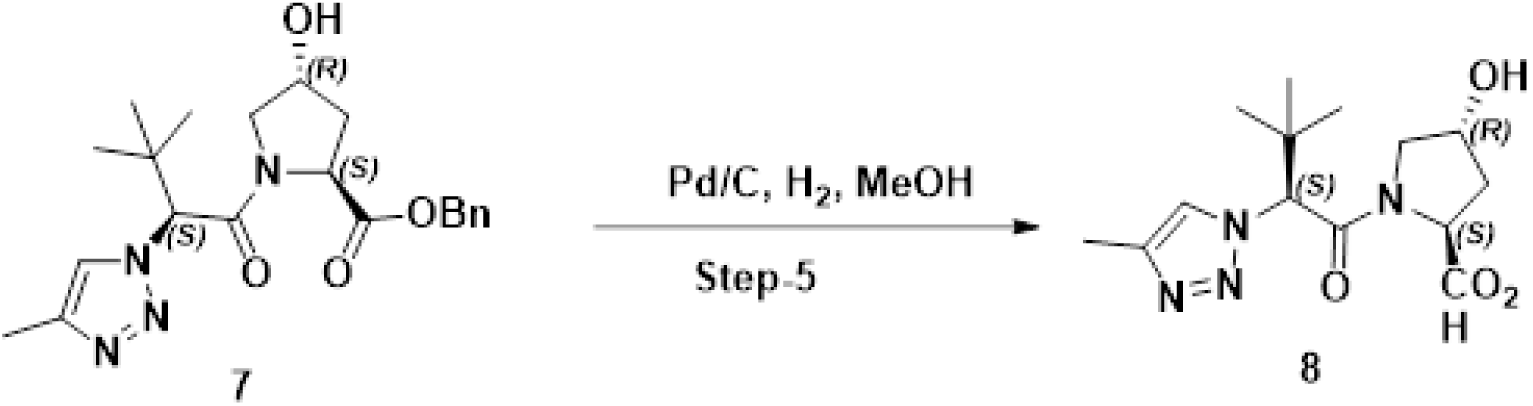

To a solution of benzyl (2*S*,4*R*)-1-((*S*)-3,3-dimethyl-2-(4-methyl-1*H*-1,2,3-triazol-1-yl)butanoyl)-4-hydroxypyrrolidine-2-carboxylate (**7**, 5.2 g, 12.98 mmol, 1.0 equiv) in methanol (30 mL) was added Pd/C (10% on wet basis, 0.650 g, 0.611 mmol, 0.5 equiv) at 25 °C. The resulting mixture was degassed and purged with hydrogen gas and stirred under hydrogen atmosphere (50 psi) at 25 °C for 2 h. The reaction mass was filtered through a celite bed and washed with methanol (100 mL). The filtrate was concentrated under reduced pressure. The crude material was triturated with acetonitrile (15 mL), filtered and dried under vacuum to give (2*S*,4*R*)-1-((*S*)-3,3-dimethyl-2-(4-methyl-1*H*-1,2,3-triazol-1-yl)butanoyl)-4-hydroxypyrrolidine-2-carboxylic acid (**8**, 2.05 g, 6.30 mmol, 48.5 % yield) as a white solid.

**LCMS (ESI, Positive ion) *m/z*:** 311.34 (M+H)^+^

**^1^H NMR (400 MHz, DMSO-*d₆*:)** δ 12.60 (br s, 1H), 7.99 (s, 1H), 5.40 (s, 1H), 5.19 (d, *J* = 3.5 Hz, 1H), 4.34 (br s, 1H), 4.25 (t, *J* = 8.4 Hz, 1H), 3.73 (dd, *J* = 10.8, 3.8 Hz, 1H), 3.61 (d, *J* = 11.0 Hz, 1H), 2.24 (s, 3H), 2.15 – 2.08 (m, 1H), 1.95 – 1.87 (m, 1H), 0.99 (s, 9H).

**Chiral Purity:** 97.1 % ee

**Chiral Purity Method:**

Column/dimensions: Chiralcel OX-H (4.6 x 250) mm,5μ% CO2 :75%, % Co solvent :25% (0.5% DEA in MeOH), Flow: 3mL/min, Back Pressure :1500 psi, Temperature :30°C

### Experimental procedure for dCASP2-1, dCASP2-2, dCASP2-3, dCASP2-4

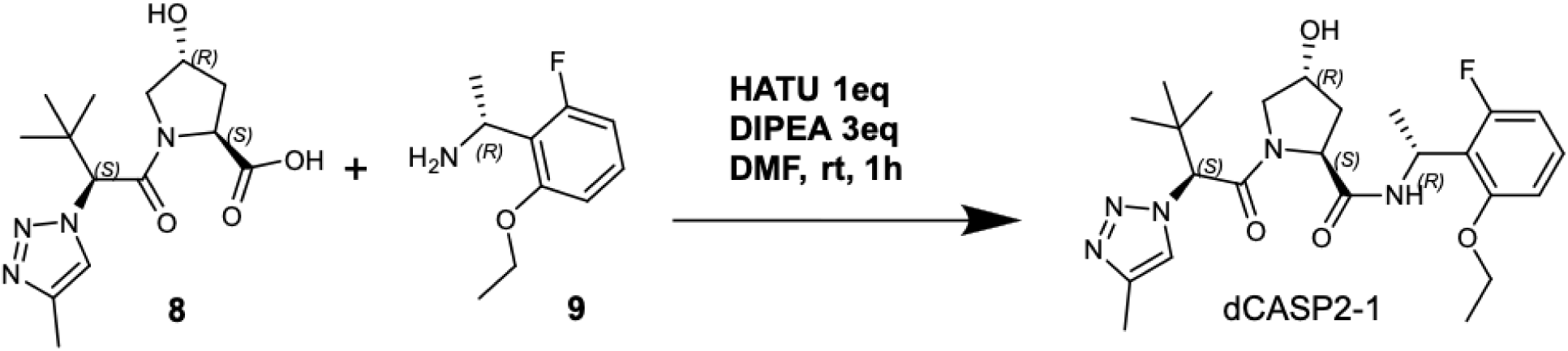

### Synthesis of (2S,4R)-1-((S)-3,3-dimethyl-2-(4-methyl-1H-1,2,3-triazol-1-yl)butanoyl)-N-((R)-1-(2-ethoxy-6-fluorophenyl)ethyl)-4-hydroxypyrrolidine-2-carboxamide (dCASP2-1)

To a 4mL dram vial containing (R)-1-(2-ethoxy-6-fluorophenyl)ethan-1-amine (**9**, 7.3 mg, 40 μmol) of were added (2S,4R)-1-((S)-3,3-dimethyl-2-(4-methyl-1H-1,2,3-triazol-1-yl)butanoyl)-4-hydroxypyrrolidine-2-carboxylic acid (**8**, 12.41 mg, 40.0 µmol), HATU (15.21 mg, 40.0 µmol), and N-ethyl-N-isopropylpropan-2-amine (DIPEA, 15.51 mg, 120 µmol).

The reaction mixture was dissolved in 600uL of DMF and stirred for one hour at room temperature. The reaction was purified by HPLC using an XSelect C18, 19 x 100 mm, 5 um, with acetonitrile w/ 0.1% formic acid in water w/ 0.1% formic acid. MS mode: ESI+. The fractions were concentrated under reduced pressure to afford the desired product (**dCASP2-1**, Yield: 63%, 11.9 mg).

**LCMS (ESI, Positive ion) *m/z*:** 476.2 (M+H)^+^ rt 0.93 min

**¹H NMR (400 MHz, DMSO-*d₆*)** δ ppm 7.89 (br d, J = 8.1 Hz, 1H, NH), 7.89 (d, J = 0.7 Hz, 1H), 7.23 (td, J = 8.3, 6.8 Hz, 1H), 6.84 (d, J = 8.4 Hz, 1H), 6.75 (ddd, J = 10.1, 8.5, 0.6 Hz, 1H), 5.42 (quin, J = 7.2 Hz, 1H), 5.36 (s, 1H), 4.45 (s, 1H), 4.32–4.11 (m, 3H), 3.72 (s, 1H), 3.49 (m, 2H), 2.22 (d, J = 0.6 Hz, 3H), 1.97 (m, 2H), 1.39 (t, J = 7.0 Hz, 3H), 0.86 (s, 9H).

**13C NMR (151 MHz, DMSO- *d₆*)** δ 169.75, 165.82, 160.57 (d, *J* = 243.2 Hz, 1C), 157.44 (d, *J* = 7.7 Hz, 1C), 141.42, 128.83, 122.20, 118.16 (d, *J* = 15.4 Hz, 1C), 108.56 (d, *J* = 2.2 Hz, 1C), 107.85 (d, *J* = 23.1 Hz, 1C), 68.68, 67.27, 64.38, 58.59, 56.48, 37.30, 35.93, 26.14 (s, 3C), 20.11, 14.57, 10.48

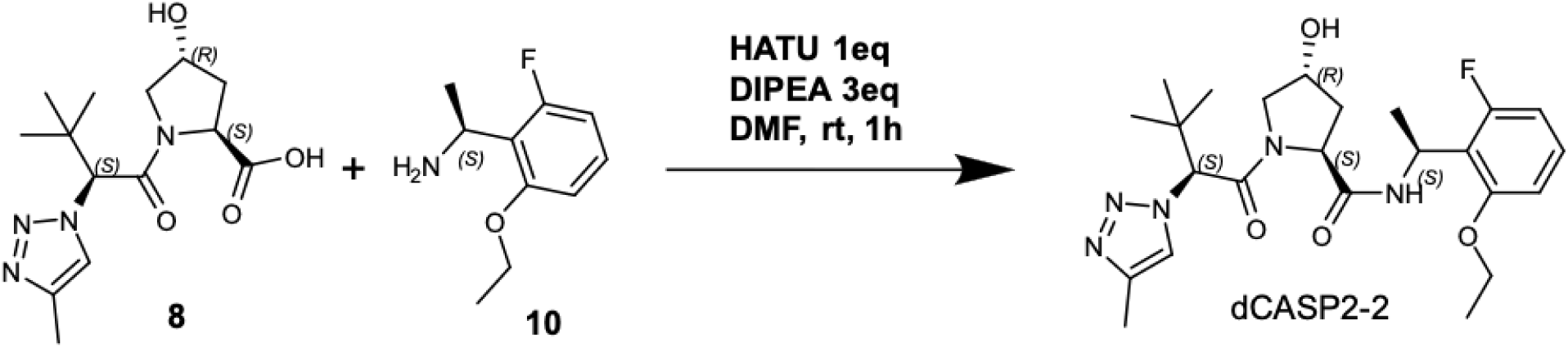

### Synthesis of (2S,4R)-1-((S)-3,3-dimethyl-2-(4-methyl-1H-1,2,3-triazol-1-yl)butanoyl)-N-((S)-1-(2-ethoxy-6-fluorophenyl)ethyl)-4-hydroxypyrrolidine-2-carboxamide (dCASP2-2)

To a 4mL dram vial containing (S)-1-(2-ethoxy-6-fluorophenyl)ethan-1-amine (**10**, 7.3 mg, 40umol) were added (2S,4R)-1-((S)-3,3-dimethyl-2-(4-methyl-1H-1,2,3-triazol-1-yl)butanoyl)-4-hydroxypyrrolidine-2-carboxylic acid (**8**, 12.41 mg, 40.0 µmol), HATU (15.21 mg, 40.0 µmol), and N-ethyl-N-isopropylpropan-2-amine (DIPEA, 15.51 mg, 120 µmol). The reaction mixture was dissolved in 600uL of DMF and stirred for one hour at room temperature. The reaction was purified by HPLC using an XSelect C18, 19 x 100 mm, 5 um, with acetonitrile w/ 0.1% formic acid in water w/ 0.1% formic acid. MS mode: ESI+. The fractions were concentrated under reduced pressure to afford the desired product.

(**dCASP2-2**, Yield: 69%, 13.1 mg).

**LCMS (ESI, Positive ion) *m/z*:** 476.2 (M+H)^+^ rt 1.11 min

**¹H NMR (400 MHz, DMSO- *d₆*)** δ ppm 8.09 (d, J = 8.1 Hz, 1H, NH), 8.01 (d, J = 0.6 Hz, 1H), 6.93 (dd, J = 8.3, 6.6 Hz, 1H), 6.60 (dd, J = 10.3, 8.6 Hz, 1H), 5.41 (s, 1H), 5.29 (quin, J = 7.3 Hz, 1H), 4.46 (t, J = 8.1 Hz, 1H), 4.25 (tt, J = 4.3, 2.3 Hz, 1H), 4.20 (t, J = 5.2 Hz, 2H), 3.67–3.55 (m, 2H), 2.69 (br t, J = 6.5 Hz, 2H), 2.24 (s, 3H), 1.93–1.74 (m, 2H), 1.35 (d, J = 7.0 Hz, 3H), 0.98 (s, 9H).

**13C NMR (151 MHz, DMSO- *d₆*)** δ 169.65, 165.76, 160.50 (d, *J* = 242.1 Hz, 1C), 157.20 (d, *J* = 8.8 Hz, 1C), 141.46, 128.54 (d, *J* = 11.0 Hz, 1C), 122.29, 118.79 (d, *J* = 14.3 Hz, 1C), 108.27, 107.80 (d, *J* = 23.1 Hz, 1C), 68.63, 67.31, 64.16, 58.52, 56.65, 37.23, 36.08, 26.27 (s, 3C), 19.87, 14.59, 10.50

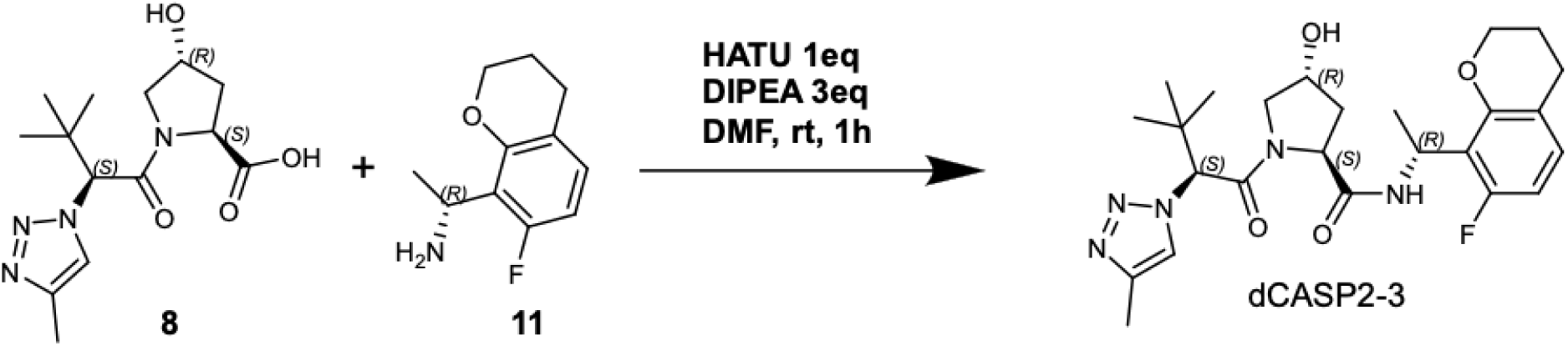

### Synthesis of (2S,4R)-1-((S)-3,3-dimethyl-2-(4-methyl-1H-1,2,3-triazol-1-yl)butanoyl)-N-((R)-1-(7-fluorochroman-8-yl)ethyl)-4-hydroxypyrrolidine-2-carboxamide (dCASP2-*3*)

To a 4mL dram vial containing (R)-1-(7-fluorochroman-8-yl)ethan-1-amine (**11**, 7.8 mg, 40umol), were added (2S,4R)-1-((S)-3,3-dimethyl-2-(4-methyl-1H-1,2,3-triazol-1-yl)butanoyl)-4-hydroxypyrrolidine-2-carboxylic acid (**8**, 12.41 mg, 40.0 µmol), HATU (15.21 mg, 40.0 µmol), and N-ethyl-N-isopropylpropan-2-amine (DIPEA, 15.51 mg, 120 µmol). The reaction mixture was dissolved in 600uL of DMF and stirred for one hour at room temperature. The crude mixture was purified by HPLC using an XSelect C18, 19 x 100 mm, 5 um, with acetonitrile w/ 0.1% formic acid in water w/ 0.1% formic acid. MS mode: ESI+. The fractions were concentrated under reduced pressure to afford the desired product (**dCASP2-3**, Yield: 43%, 8.4 mg).

**LCMS (ESI, Positive ion) *m/z*:** 488.0 (M+H)^+^ rt 0.93 min

**¹H NMR (400 MHz, DMSO- *d₆*):** δ 7.90 (d, J = 0.8 Hz, 1H, H-10), 7.83 (d, J = 8.5 Hz, 1H, NH-22), 6.97 (dd, J = 10.2, 8.5 Hz, 1H, H-32), 6.63 (dd, J = 10.1, 8.5 Hz, 1H, H-33), 5.38 (s, 1H, H-8), 5.36 (dq, J = 8.5, 6.8 Hz, 1H, H-23), 4.44 (t, J = 7.8 Hz, 1H, H-2), 4.31 (m, 1H, H-4), 4.26–4.22 (ddd, J = 10.7, 6.6, 3.8 Hz, 2H, H-30), 3.73 (dd, J = 10.7, 4.4 Hz, 1H, H-5), 3.48 (dd, J = 10.8, 0.9 Hz, 1H, H-5), 2.25–1.93 (m, 4H, H-3, H-29), 2.22 (s, 3H, H-12), 1.32 (d, J = 6.8 Hz, 3H, H-24), 0.88 (s, 9H, H-14–18).

**13C NMR (151 MHz, DMSO- *d₆*)** δ 169.80, 165.86, 158.66 (d, *J* = 241.0 Hz, 1C), 152.93 (d, *J* = 8.3 Hz, 1C), 141.42, 128.93 (d, *J* = 10.2 Hz, 1C), 122.19, 118.43 (d, *J* = 3.0 Hz, 1C), 117.02 (d, *J* = 16.5 Hz, 1C), 106.81 (d, *J* = 23.1 Hz, 1C), 68.61, 67.25, 66.33, 58.72, 56.44, 37.38, 35.92, 26.16 (s, 3C), 23.93, 21.31, 20.26, 10.48

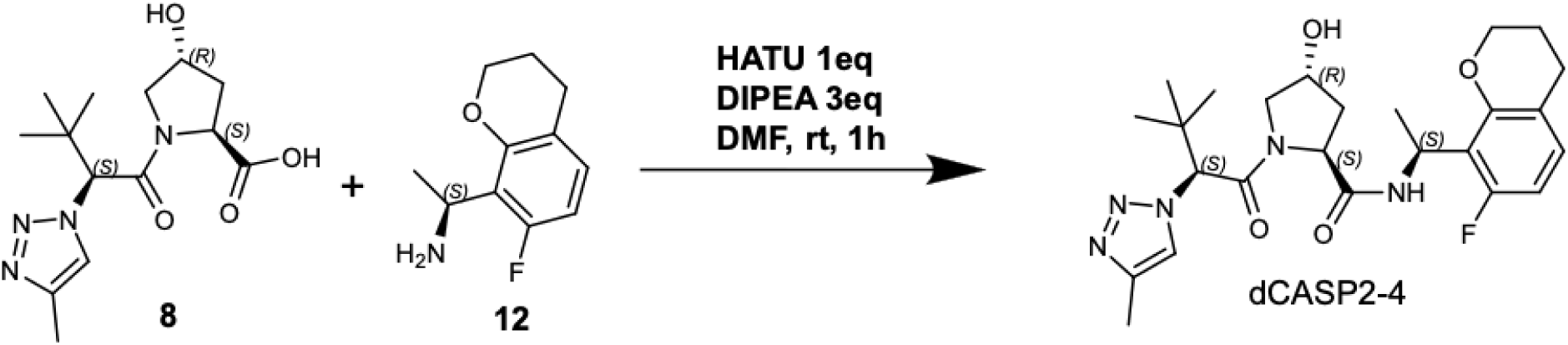

### Synthesis of (2S,4R)-1-((S)-3,3-dimethyl-2-(4-methyl-1H-1,2,3-triazol-1-yl)butanoyl)-N-((S)-1-(7-fluorochroman-8-yl)ethyl)-4-hydroxypyrrolidine-2-carboxamide (dCASP2-4)

To a 4mL dram vial containing (S)-1-(7-fluorochroman-8-yl)ethan-1-amine (**12**, 7.8 mg, 40umol), were added (2S,4R)-1-((S)-3,3-dimethyl-2-(4-methyl-1H-1,2,3-triazol-1-yl)butanoyl)-4-hydroxypyrrolidine-2-carboxylic acid (**8**, 12.41 mg, 40.0 µmol), HATU (15.21 mg, 40.0 µmol), and N-ethyl-N-isopropylpropan-2-amine (DIPEA, 15.51 mg, 120 µmol). The reaction mixture was dissolved in 600uL of DMF and stirred for one hour at room temperature. The crude mixture was purified by HPLC using an XSelect C18, 19 x 100 mm, 5 um, with acetonitrile w/ 0.1% formic acid in water w/ 0.1% formic acid. MS mode: ESI+. The fractions were concentrated under reduced pressure to afford the desired product (**dCASP2-4**, Yield: 71%, 13.6 mg).

**LCMS (ESI, Positive ion) *m/z*:** 488.2 (M+H)^+^ rt 0.91 min

**¹H NMR (400 MHz, DMSO- *d₆*)** δ ppm 8.09 (d, J = 8.1 Hz, 1H, NH), 8.01 (d, J = 0.6 Hz, 1H), 6.93 (dd, J = 8.3, 6.6 Hz, 1H), 6.60 (dd, J = 10.3, 8.6 Hz, 1H), 5.41 (s, 1H), 5.29 (quin, J = 7.3 Hz, 1H), 4.46 (t, J = 8.1 Hz, 1H), 4.25 (tt, J = 4.3, 2.3 Hz, 1H), 4.20 (t, J = 5.2 Hz, 2H), 3.67 (dd, J = 10.8, 4.2 Hz, 1H), 3.55 (dt, J = 11.0, 1.8 Hz, 1H), 2.69 (br t, J = 6.5 Hz, 2H), 2.24 (s, 3H), 1.93 (m, 1H), 1.74 (ddd, J = 12.9, 8.2, 4.8 Hz, 1H), 1.35 (d, J = 7.0 Hz, 3H), 0.98 (s, 9H).

**13C NMR (151 MHz, DMSO- *d₆*)** δ 169.69, 165.69, 158.67 (d, *J* = 241.0 Hz, 1C), 152.74 (d, *J* = 7.7 Hz, 1C), 141.45, 128.69 (d, *J* = 11.0 Hz, 1C), 122.35, 118.25 (d, *J* = 3.3 Hz, 1C), 117.83 (d, *J* = 15.4 Hz, 1C), 106.83 (d, *J* = 22.0 Hz, 1C), 68.66, 67.34, 66.27, 58.49, 56.70, 37.28, 36.10, 26.28

(s, 3C), 23.89, 21.38, 19.98, 10.49

## Acknowledgements

We thank Kimberly Garcia from the Research Material Management group of Amgen Research for the help with sample and compound management. In addition, we appreciate the discussion with Kate Ashton from the Medicinal Chemistry, Emily Cook, Kusal Samarasinghe from the Induced-Proximity Platform, Xiaomin Chen, Sudipa Ghimire-Rijal, Hui-Ting Chou, Angela Lei from the Structural Biology and Ann Shim from the Discovery Protein Science groups of Amgen Research for their insightful discussions.

## Author contributions

J.M., A.S., and J.H. designed the study and interpreted the results; J.M. designed the compounds; A.G. synthesized the molecular glue compounds; J.H. performed cellular characterization of compounds; K.S. performed JESS quantification analyses; A.I., K.Chen, and K.Cho performed biochemical characterization; S.L. generated the TurboID cell line; W.D. performed proteomics analyses; S.O. carried out computational modeling of the ternary complex and performed molecular dynamics simulations; S.J.B., S.Z., K.H.C., W.B., S.V., O.B., B.Z. contributed reagents, materials, and analysis tools; P.R.P. supervised the study; J.M., A.S., and J.H. wrote the manuscript.

## Declaration of competing interests

Financial support for this research was provided by Amgen. All authors are employees of Amgen and may hold stock or stock options in the company. The authors declare no other competing interests.

## Availability of data sharing

All data associated with this study are present in the paper or Supplementary materials. Requests for materials should be sent to the corresponding authors and material transfer agreements are required.

## Notes

### Competing Interest Statement

The authors have declared no competing interest.

### Summary of Updates

No major revisions were made. One co-author who was inadvertently omitted from the author list has been added, and several synthetic schemes in the Methods section that were not displayed correctly in the PDF version have been corrected.

## Reference

1 Yoon, H., Rutter, J. C., Li, Y.-D. & Ebert, B. L. Induced protein degradation for therapeutics: past, present, and future. The Journal of Clinical Investigation 134 (2024). 10.1172/JCI175265

2 Zhong, G., Chang, X., Xie, W. & Zhou, X. Targeted protein degradation: advances in drug discovery and clinical practice. Signal Transduction and Targeted Therapy 9, 308 (2024). 10.1038/s41392-024-02004-x

3 Békés, M., Langley, D. R. & Crews, C. M. PROTAC targeted protein degraders: the past is prologue. Nature Reviews Drug Discovery 21, 181–200 (2022). 10.1038/s41573-021-00371-6

4 Burslem, G. M. & Crews, C. M. Proteolysis-Targeting Chimeras as Therapeutics and Tools for Biological Discovery. Cell 181, 102–114 (2020). 10.1016/j.cell.2019.11.031

5 Konstantinidou, M. & Arkin, M. R. Molecular glues for protein-protein interactions: Progressing toward a new dream. Cell Chemical Biology 31, 1064–1088 (2024). 10.1016/j.chembiol.2024.04.002

6 Oleinikovas, V., Gainza, P., Ryckmans, T., Fasching, B. & Thomä, N. H. From Thalidomide to Rational Molecular Glue Design for Targeted Protein Degradation. Annual Review of Pharmacology and Toxicology 64, 291–312 (2024). 10.1146/annurev-pharmtox-022123-104147

7 Schreiber, S. L. Molecular glues and bifunctional compounds: Therapeutic modalities based on induced proximity. Cell Chemical Biology 31, 1050–1063 (2024). 10.1016/j.chembiol.2024.05.004

8 Sasso, J. M. et al. Molecular Glues: The Adhesive Connecting Targeted Protein Degradation to the Clinic. Biochemistry 62, 601–623 (2023). 10.1021/acs.biochem.2c00245

9 Ito, T. et al. Identification of a Primary Target of Thalidomide Teratogenicity. Science 327, 1345–1350 (2010). doi:10.1126/science.1177319

10 Krönke, J. et al. Lenalidomide Causes Selective Degradation of IKZF1 and IKZF3 in Multiple Myeloma Cells. Science 343, 301–305 (2014). doi:10.1126/science.1244851

11 Sievers, Q. L. et al. Defining the human C2H2 zinc finger degrome targeted by thalidomide analogs through CRBN. Science 362, eaat0572 (2018). doi:10.1126/science.aat0572

12 Petzold, G., Fischer, E. S. & Thomä, N. H. Structural basis of lenalidomide-induced CK1α degradation by the CRL4CRBN ubiquitin ligase. Nature 532, 127–130 (2016). 10.1038/nature16979

13 Krönke, J. et al. Lenalidomide induces ubiquitination and degradation of CK1α in del(5q) MDS. Nature 523, 183–188 (2015). 10.1038/nature14610

14 Lu, G. et al. The Myeloma Drug Lenalidomide Promotes the Cereblon-Dependent Destruction of Ikaros Proteins. Science 343, 305–309 (2014). doi:10.1126/science.1244917

15 Rajkumar, S. V. et al. Combination therapy with lenalidomide plus dexamethasone (Rev/Dex) for newly diagnosed myeloma. Blood 106, 4050–4053 (2005). 10.1182/blood-2005-07-2817

16 Hansen, J. D. et al. Discovery of CRBN E3 Ligase Modulator CC-92480 for the Treatment of Relapsed and Refractory Multiple Myeloma. Journal of Medicinal Chemistry 63, 6648–6676 (2020). 10.1021/acs.jmedchem.9b01928

17 Baek, K. et al. Unveiling the hidden interactome of CRBN molecular glues. Nature Communications 16, 6831 (2025). 10.1038/s41467-025-62099-w

18 Petzold, G. et al. Mining the CRBN target space redefines rules for molecular glue–induced neosubstrate recognition. Science 389, eadt6736 (2025). doi:10.1126/science.adt6736

19 Steger, M. et al. Unbiased mapping of cereblon neosubstrate landscape by high-throughput proteomics. Nature Communications 16, 7773 (2025). 10.1038/s41467-025-62829-0

20 Annunziato, S. et al. Cereblon induces G3BP2 neosubstrate degradation using molecular surface mimicry. Nature Structural & Molecular Biology 33, 479–487 (2026). 10.1038/s41594-025-01738-8

21 Ivan, M. et al. HIFalpha Targeted for VHL-Mediated Destruction by Proline Hydroxylation: Implications for O2 Sensing. Science 292, 464–468 (2001). doi:10.1126/science.1059817

22 Frost, J. et al. Potent and selective chemical probe of hypoxic signalling downstream of HIF-α hydroxylation via VHL inhibition. Nature Communications 7, 13312 (2016). 10.1038/ncomms13312

23 Galdeano, C. et al. Structure-Guided Design and Optimization of Small Molecules Targeting the Protein–Protein Interaction between the von Hippel–Lindau (VHL) E3 Ubiquitin Ligase and the Hypoxia Inducible Factor (HIF) Alpha Subunit with in Vitro Nanomolar Affinities. Journal of Medicinal Chemistry 57, 8657–8663 (2014). 10.1021/jm5011258

24 Diehl, C. J. & Ciulli, A. Discovery of small molecule ligands for the von Hippel-Lindau (VHL) E3 ligase and their use as inhibitors and PROTAC degraders. Chemical Society Reviews 51, 8216–8257 (2022). 10.1039/D2CS00387B

25 Bondeson, D. P. et al. Catalytic in vivo protein knockdown by small-molecule PROTACs. Nature Chemical Biology 11, 611–617 (2015). 10.1038/nchembio.1858

26 Zengerle, M., Chan, K.-H. & Ciulli, A. Selective Small Molecule Induced Degradation of the BET Bromodomain Protein BRD4. ACS Chemical Biology 10, 1770–1777 (2015). 10.1021/acschembio.5b00216

27 Khan, S. et al. A selective BCL-XL PROTAC degrader achieves safe and potent antitumor activity. Nature Medicine 25, 1938–1947 (2019). 10.1038/s41591-019-0668-z

28 Zhao, H.-Y. et al. Discovery of Potent PROTACs Targeting EGFR Mutants through the Optimization of Covalent EGFR Ligands. Journal of Medicinal Chemistry 65, 4709–4726 (2022). 10.1021/acs.jmedchem.1c01827

29 Kofink, C. et al. A selective and orally bioavailable VHL-recruiting PROTAC achieves SMARCA2 degradation in vivo. Nature Communications 13, 5969 (2022). 10.1038/s41467-022-33430-6

30 Tutter, A. et al. A small-molecule VHL molecular glue degrader for cysteine dioxygenase 1. Nature Chemical Biology (2025). 10.1038/s41589-025-01936-x

31 Bushman, J. W. et al. Discovery of a VHL molecular glue degrader of GEMIN3 by Picowell RNA-seq. bioRxiv, 2025.2003.2019.644003 (2025). 10.1101/2025.03.19.644003

32 Kopeina, G. S. & Zhivotovsky, B. Caspase-2 as a master regulator of genomic stability. Trends in Cell Biology 31, 712–720 (2021). 10.1016/j.tcb.2021.03.002

33 Sladky, V. C. & Villunger, A. Uncovering the PIDDosome and caspase-2 as regulators of organogenesis and cellular differentiation. Cell Death & Differentiation 27, 2037–2047 (2020). 10.1038/s41418-020-0556-6

34 Ando, K. et al. NPM1 directs PIDDosome-dependent caspase-2 activation in the nucleolus. Journal of Cell Biology 216, 1795–1810 (2017). 10.1083/jcb.201608095

35 Vakifahmetoglu-Norberg, H. & Zhivotovsky, B. The unpredictable caspase-2: what can it do? Trends in Cell Biology 20, 150–159 (2010). 10.1016/j.tcb.2009.12.006

36 Kumar, S. Caspase 2 in apoptosis, the DNA damage response and tumour suppression: enigma no more? Nature Reviews Cancer 9, 897–903 (2009). 10.1038/nrc2745

37 Shashikadze, B. et al. Integrated proteomic screening reveals design principles of CRBN molecular glue degraders. bioRxiv, 2026.2003.2008.710269 (2026). 10.64898/2026.03.08.710269

38 Friesner, R. A. et al. Glide: A New Approach for Rapid, Accurate Docking and Scoring. 1. Method and Assessment of Docking Accuracy. Journal of Medicinal Chemistry 47, 1739-1749 (2004). 10.1021/jm0306430

39 Stebbins, C. E., Kaelin, W. G. & Pavletich, N. P. Structure of the VHL-ElonginC-ElonginB Complex: Implications for VHL Tumor Suppressor Function. Science 284, 455–461 (1999). doi:10.1126/science.284.5413.455

40. Kozakov, D., Brenke, R., Comeau, S. R. & Vajda, S. PIPER: An FFT-based protein docking program with pairwise potentials. Proteins: Structure, Function, and Bioinformatics 65, 392-406 (2006). 10.1002/prot.21117

41 Wang, L. et al. Accurate and Reliable Prediction of Relative Ligand Binding Potency in Prospective Drug Discovery by Way of a Modern Free-Energy Calculation Protocol and Force Field. Journal of the American Chemical Society 137, 2695–2703 (2015). 10.1021/ja512751q

