## Supplementary Information for "Reprogramming VHL with molecular glues enables selective degradation of caspase-2"

### Supplementary Figures

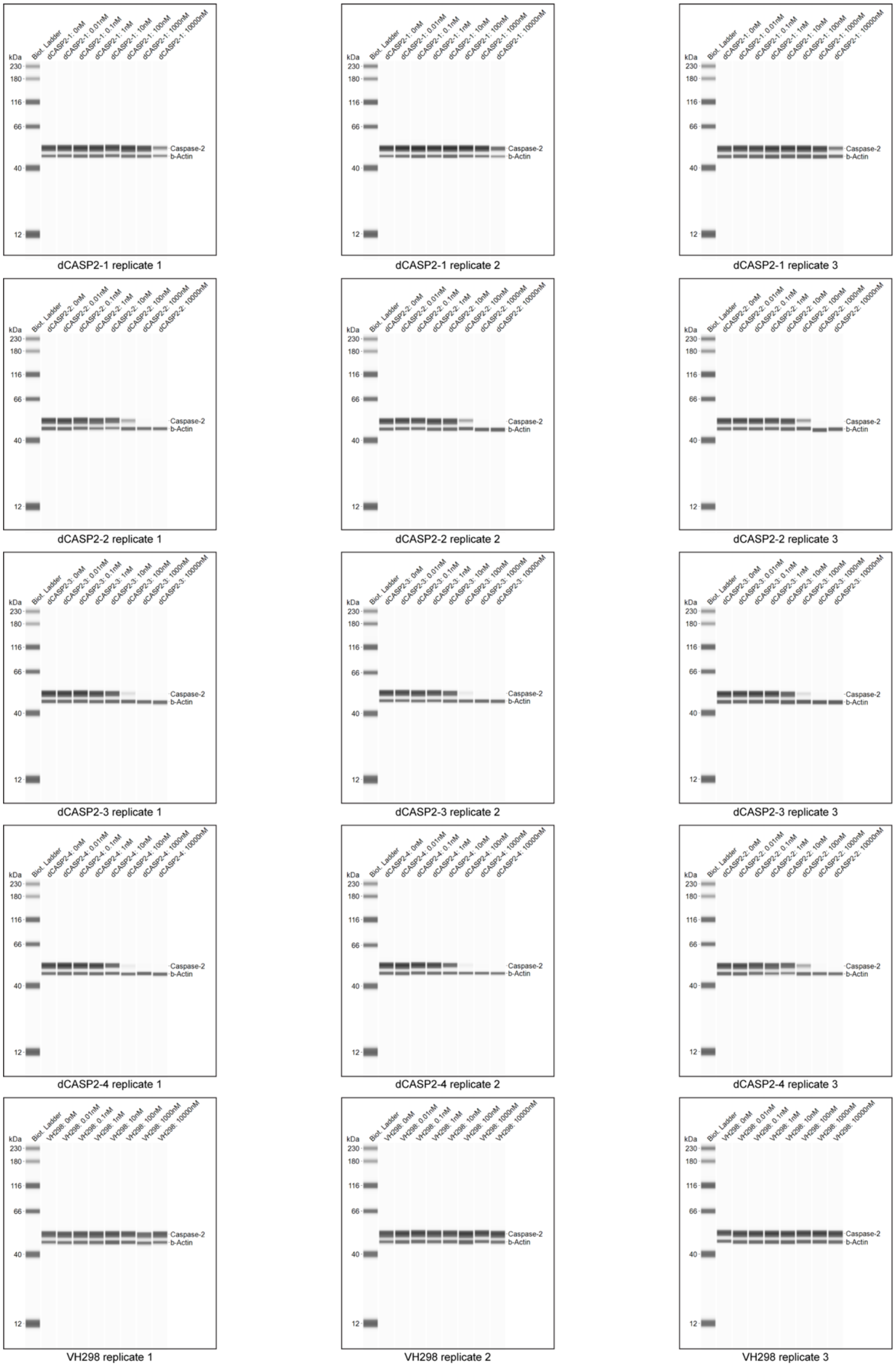


**Supplementary Figure S1. JESS Image for quantification of caspase-2 levels.** Jurkat cell lysates were collected after 24-hour treatment with increasing concentrations of caspase-2 molecular glue degraders and negative control VH298. JESS quantification was performed in three biological replicates. β-actin was used as a loading control.

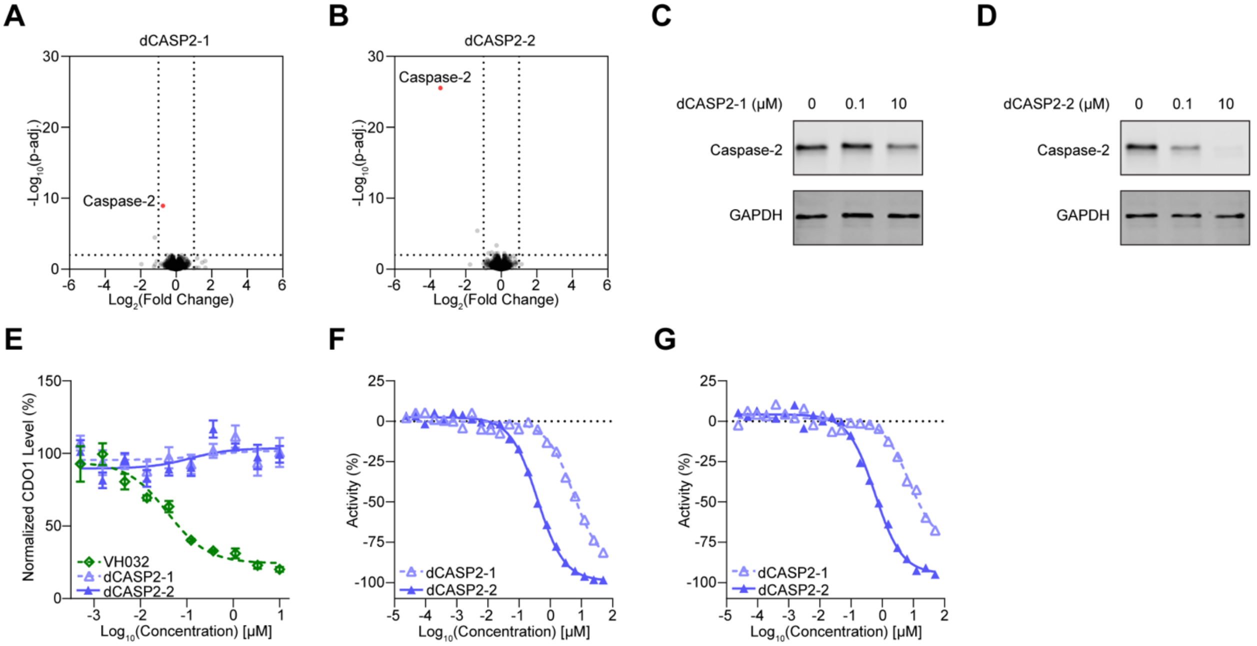


**Supplementary Figure S2. Rapid and selective caspase-2 degradation by dCASP2-1 and dCASP2-2. A-B.** dCASP2-1 and dCASP2-2 promote caspase-2 downregulation. Wild-type Jurkat cells were treated for 6 hours with dCASP2-1 (10 µM, **A**, n=3), dCASP2-2 (10 µM, **B**, n=3), or DMSO (0.1%, n=3), followed by global proteomics analysis. Dashed lines indicate significance and enrichment cutoffs of adjusted p-value < 0.01 and $log_2$ fold-change (relative to DMSO) < –1. Caspase-2 is highlighted in red. **C-D.** dCASP2-1 and dCASP2-2 rapidly degrade caspase-2. Jurkat cells were treated for 6 hours with increasing concentrations of dCASP2-1 (**C**) or dCASP2-2 (**D**). Endogenous caspase-2 was analyzed by Western blot, with GAPDH probed as a loading control. **E.** dCASP2-1 and dCASP2-2 are inactive in a CDO1 degradation assay. HiBiT assays were used to measure CDO1 levels after 24-hour treatment with dCASP2-1 or dCASP2-2. VH032 was included as a positive control (n=3). **F-G.** dCASP2-1 and dCASP2-2 bind to VHL. VBC NanoBRET VHL target engagement assays were performed in permeabilized (**F**) and live cells (**G**). Decreased % activity indicates increased VHL binding (n=1).

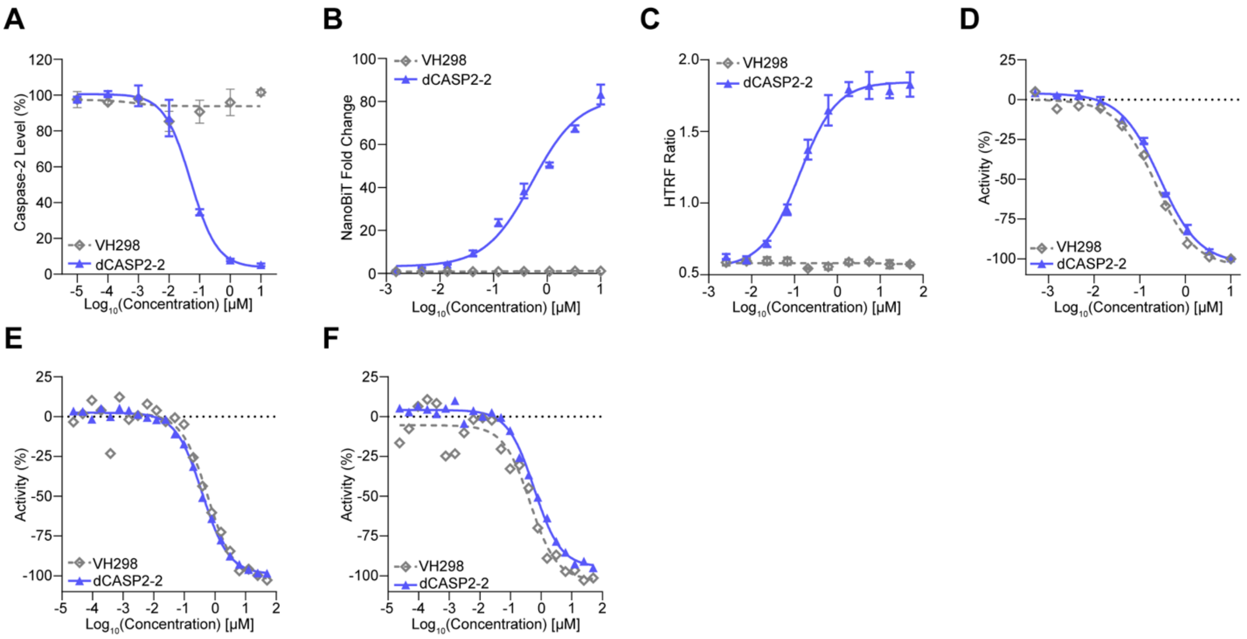


**Supplementary Figure S3. VH298 is not a caspase-2 degrader (dCASP2-2 was used as positive control). A.** VH298 does not degrade caspase-2 in a dose-dependent manner. Caspase-2 levels were quantified by JESS following 24-hour VH298 treatment at increasing concentrations in Jurkat cells (n=3). **B.** VH298 does not induce caspase-2/MGD/VHL ternary complex formation in cells. HEK293T cells were transiently co-transfected with VHL-LgBiT and SmBiT-caspase-2 vectors, then co-treated with MLN4924 and caspase-2 degraders. Increased luminescence reflects compound-induced ternary complex formation (n=3). **C.** VH298 does not induce *in vitro* caspase-2/MGD/VHL ternary complex formation. HTRF assays were performed with purified caspase-2 and VBC complex in the presence of VH298. Higher HTRF signal indicates ternary complex formation *in vitro* (n=3). **D-F.** VH298 binds to VHL. VBC AlphaScreen competition assays were used to determine binary binding of VH298 to VHL *in vitro* (**D**). VBC NanoBRET VHL target engagement assays were also performed in permeabilized (**E**) and live cells (**F**). In all assays, decreased % activity indicates stronger VHL engagement (n=1).

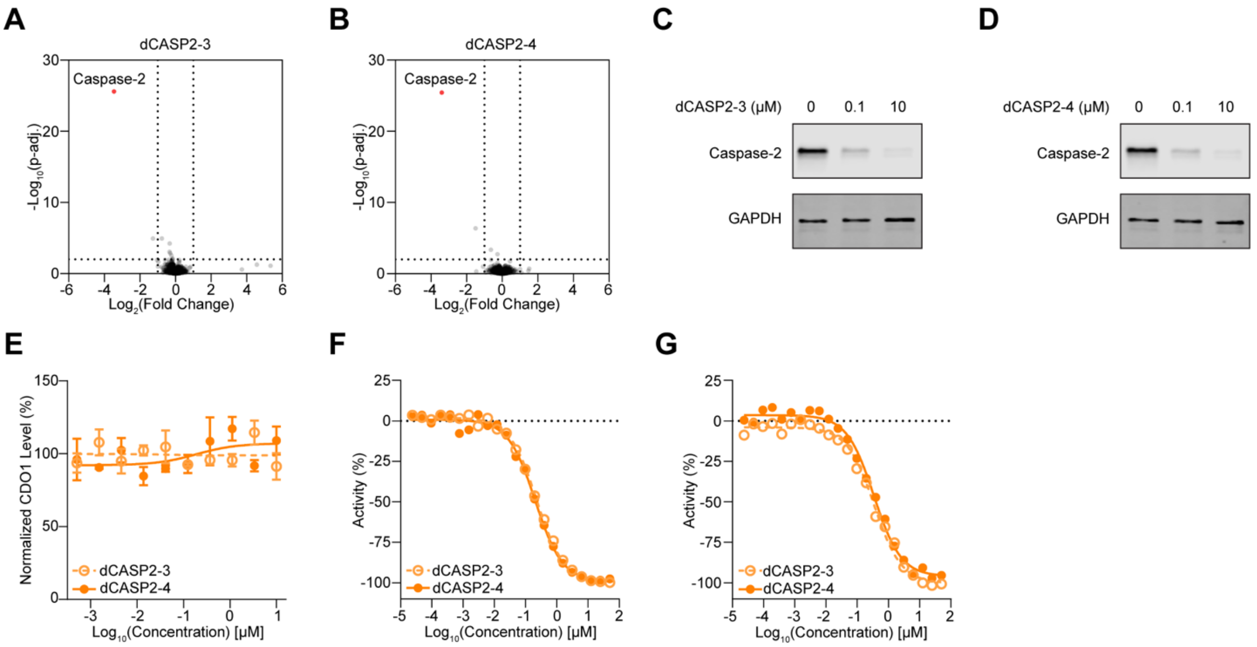


**Supplementary Figure S4. Rapid and selective caspase-2 degradation by dCASP2-3 and dCASP2-4. A-B.** dCASP2-3 and dCASP2-4 promote caspase-2 downregulation. Wild-type Jurkat cells were treated for 6 hours with dCASP2-3 (10 μM, **A**, n=3), dCASP2-4 (10 μM, **B**, n=3), or DMSO (0.1%, n=3), followed by global proteomics analysis. Dashed lines indicate significance and enrichment cutoffs of adjusted p-value < 0.01 and $\log_2$ fold-change (relative to DMSO) < –1. Caspase-2 is highlighted in red. **C-D.** dCASP2-3 and dCASP2-4 rapidly degrade caspase-2. Jurkat cells were treated for 6 hours with increasing concentrations of dCASP2-3 (**C**) or dCASP2-4 (**D**). Endogenous caspase-2 was analyzed by Western blot, with GAPDH probed as a loading control. **E.** dCASP2-3 and dCASP2-4 are inactive in a CDO1 degradation assay. HiBiT assays were performed to measure CDO1 levels after 24-hour treatment with dCASP2-3 or dCASP2-4 (n=3). **F-G.** dCASP2-3 and dCASP2-4 bind to VHL. VBC NanoBRET VHL target engagement assays were performed in permeabilized (**F**) and live cells (**G**). Decreased % activity indicates stronger VHL binding (n=1).

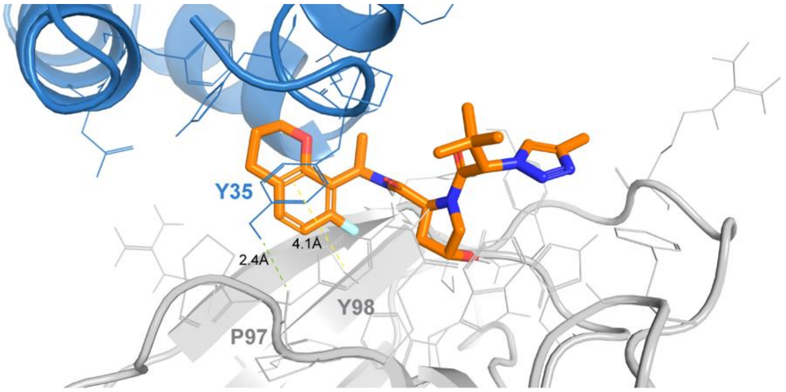


**Supplementary Figure S5. Computational 3D modeling predicts caspase-2 (H35Y)/dCASP2-3/VHL ternary complex.** Selected computational model of ternary complex of caspase-2 carrying the H35Y substitution, dCASP2-3, and VHL. The mutant caspase-2 CARD domain is shown in blue, VHL in grey, and dCASP2-3 in orange. Key interactions between mutant caspase-2 Y35 and VHL (Y98, P97) are highlighted: green line represents hydrogen bond, and yellow line shows π–π aromatic stacking.

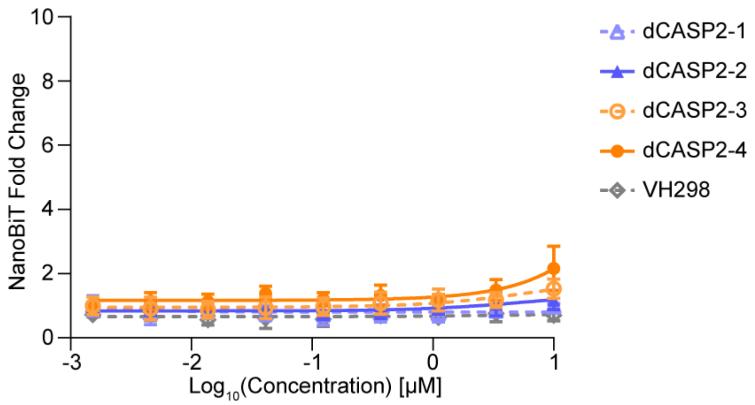


**Supplementary Figure S6. Cellular TCF assay for HIF-1α.** NanoBiT assays were performed to measure compound-induced ternary complex formation between HIF1α and VHL.
